# Contact sites between endoplasmic reticulum sheets and mitochondria regulate mitochondrial DNA replication and segregation

**DOI:** 10.1101/2021.12.09.472002

**Authors:** Hema Saranya Ilamathi, Sara Benhammouda, Amel Lounas, Khalid Al-Naemi, Justine Desrochers-Goyette, Matthew A. Lines, François J. Richard, Jackie Vogel, Marc Germain

## Abstract

Mitochondria are multi-faceted organelles crucial for cellular homeostasis that contain their own genome. Mitochondrial DNA (mtDNA) replication is a spatially regulated process essential for the maintenance of mitochondrial function, its defect causing mitochondrial diseases. mtDNA replication occurs at endoplasmic reticulum (ER)-mitochondria contact sites and is affected by mitochondrial dynamics: the absence of mitochondrial fusion is associated with mtDNA depletion whereas loss of mitochondrial fission causes the aggregation of mtDNA within abnormal structures termed mitobulbs. Here, we show that contact sites between mitochondria and ER sheets, the ER structure associated with protein synthesis, regulate mtDNA replication and distribution within mitochondrial networks. DRP1 loss or mutation leads to modified ER sheets and alters the interaction between ER sheets and mitochondria, disrupting RRBP1-SYNJ2BP interaction. Importantly, mtDNA distribution and replication were rescued by promoting ER sheets-mitochondria contact sites. Our work identifies the role of ER sheet-mitochondria contact sites in regulating mtDNA replication and distribution.

## Introduction

Mitochondria are dynamic organelles regulating an array of cellular processes including energy production, cellular metabolism, apoptosis, calcium signaling, ROS signaling, cellular differentiation, and immune response against pathogens ^1–6^. These mitochondrial functions are regulated by mitochondrial dynamics, the processes of mitochondrial fusion and fission. Mitochondrial fission requires DRP1 (Dynamin-Related Protein 1), which is recruited from the cytosol to fission sites ^7,8^. Fusion is regulated by Mitofusins 1 and 2 (MFN1 & 2) and Optic Atrophy Protein 1 (OPA1), present on the mitochondrial outer and inner membranes, respectively ^9,10^. Mitochondrial dynamics are essential for mitochondrial DNA (mtDNA) maintenance, with defects in mitochondrial fusion impairing mtDNA integrity and copy number ^11,12^. On the other hand, while defective mitochondrial fission does not generally affect overall mtDNA content, it alters the distribution of nucleoids (mtDNA-protein complexes), leading to the formation of bulb-like mitochondrial structures termed mitobulbs ^13–16^. Importantly, both mitochondrial fission and mtDNA replication are initiated at sites of contact between mitochondria and the endoplasmic reticulum (ERMCS)^17–19^, indicating a crucial role for the ER in the regulation of mitochondrial structure and function.

The ER is a complex web-like organelle that includes flattened cisternae/sheets mostly around the perinuclear region and tubulated structures towards the cellular periphery ^20^. Rough ER (rER) generally adopts a sheets-like structure that is enriched with ribosomes and regulates the production and assembly of proteins for secretion ^21^. On the other hand, smooth ER (sER) is usually present as tubulated structures that play important roles in lipid synthesis and calcium signaling, but also in mitochondrial dynamics and mtDNA replication ^17–19,21^. At the molecular level, the ER resident protein CLIMP63 (CKAP4) is essential for the stabilizing ER sheets, while reticulons (RTN) and DP1/Yop1p are essential for both ER tubule formation and the maintenance of ER sheet curvature ^22,23^. The ratio of sheet to tubule proteins regulates the balance between the two ER structures ^22^.

Specific membrane proteins present on the ER and mitochondria tether the two organelles to form ERMCS. These interorganelle tethers play key roles in the regulation of mitochondrial dynamics, mtDNA replication, calcium signaling, lipid metabolism and transfer, innate immune response, and autophagy ^17–19,24–28^. As ERMCS have mainly been reported at ER tubules, we have limited knowledge of the role of ER sheets in ERMCS formation and function. One of the few examples of ER sheet-mitochondria contact sites occurs in mouse liver cells where these ERMCS regulate lipid metabolism ^29,30^. Nevertheless, the role of ER sheets-mitochondria contact sites in mitochondrial biogenesis is poorly understood.

Here, we show that altered ER sheet-mitochondria interaction disrupts mtDNA replication and segregation. Specifically, we observe that mutation or deletion of DRP1 alters ER sheet structure and ER sheet-mitochondria interaction leading to a reduction in mtDNA replication and segregation. The deleterious effects of DRP1 mutations on mtDNA replication and distribution can be rescued by modulating ER sheet-mitochondria contact sites through manipulation of CLIMP63 and SYNJ2BP. Altogether, our results demonstrate that ER sheet-mitochondria contact sites regulate replication and distribution of mtDNA.

## Results

### Mitobulbs are clusters of mtDNA that fail to disperse

Defects or loss of fission proteins cause the appearance of large nucleoids within bulb-like mitochondrial structures termed mitobulbs ^14–16^, altering nucleoid distribution within mitochondrial networks ^14^. While this indicates an important role of mitochondrial fission in the proper distribution of nucleoids, the underlying mechanism remains unclear.

Mutation or deletion of the mitochondrial fission protein DRP1 causes the formation of mitobulbs that are primarily present in the perinuclear region of the cell ^13–16^. We observed a similar phenotype in primary fibroblasts from patients with a dominant negative mutation in the middle domain of DRP1 which is required for DRP1 oligomerization(Figure S1A) ^31^, but not in control cells (Figure 1A; mitochondria (TMRM) and DNA (picogreen), quantification of the number of mitobulbs/cell in Figure 1B). Multiple copies of replicating mtDNA were observed in individual mitobulbs when labeled with the nucleotide analog EdU (Figure 1C-D). This suggests that mtDNA can actively replicate in mitobulbs but fail to distribute along mitochondrial networks, resulting in a decrease in total nucleoid numbers ^14,15^. Consistent with this, a significant fraction of total EdU-positive nucleoids was present within mitobulbs (Figure 1E). Nevertheless, although total mtDNA levels were not decreased in DRP1 mutants (Figure 1F), they had fewer EdU-positive nucleoids compared to control cells (Figure 1G), suggesting that the loss of DRP1 function affects mtDNA replication. Altogether, our data indicate that mitochondrial fission is essential for proper mtDNA replication and distribution.

**Figure 1.**
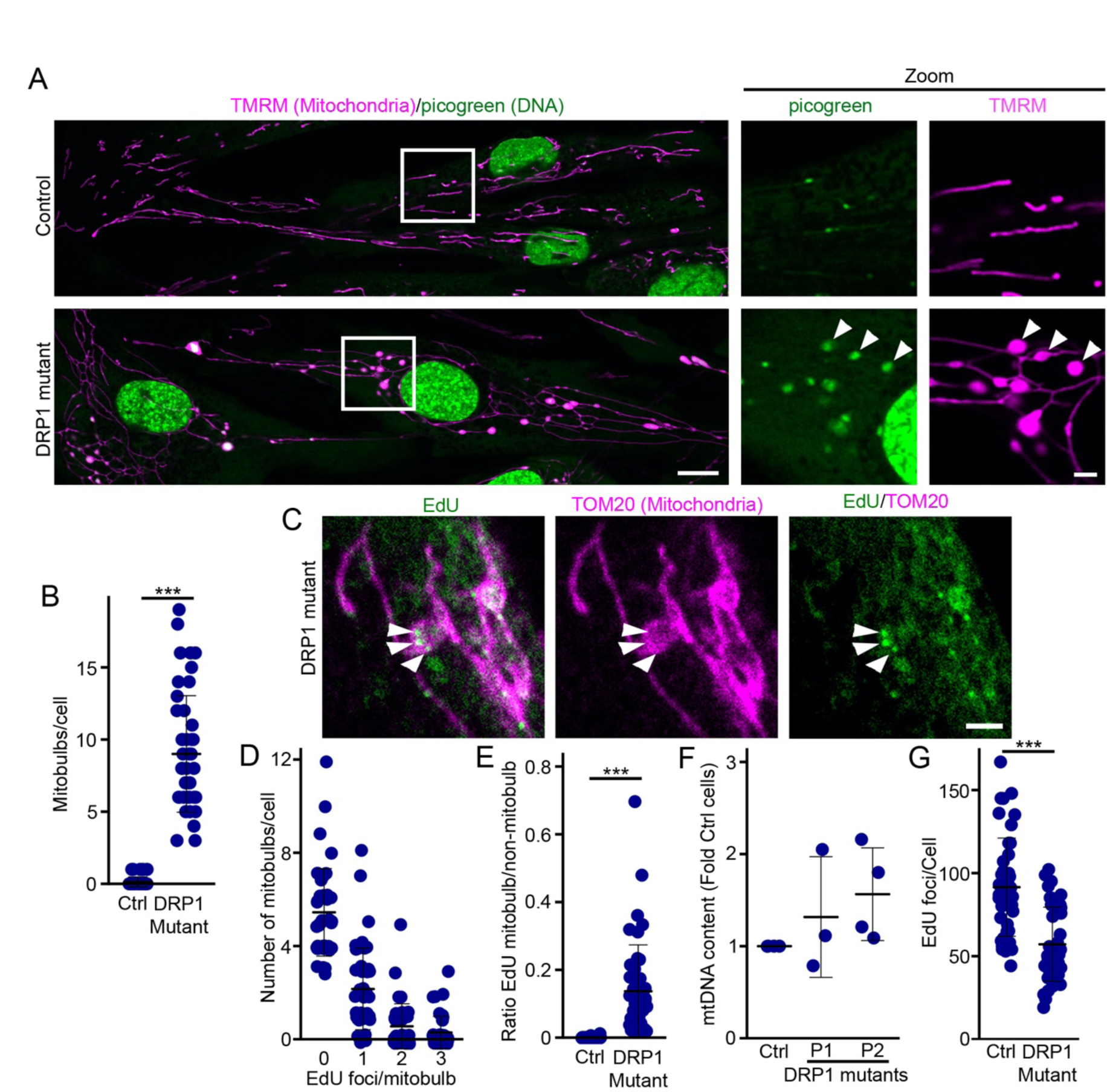
Multiple mtDNA copies accumulate within mitobulbs in DRP1 mutant fibroblasts. (A) Representative live cell images showing the mitochondrial marker TMRM (magenta) and picogreen-stained DNA (green) in control and DRP1 mutant human fibroblasts. The zoomed images show the enlarged nucleoids present in mitobulbs (arrowheads). (B) Quantification of the number of mitobulbs per cell. Each point represents an individual cell, with 45 cells quantified in 3 independent experiments. Bars show the average ± SD. (C) Representative images of mCherry-expressing DRP1 mutants labeled for EdU (green) and TOM20 (mitochondria, magenta). Arrowheads indicate EdU-positive mtDNA in mitobulbs. Scale bar 10µm. (D) Quantification of number of EdU foci/mitobulb. Each point represents an individual cell, with 45 cells quantified in 3 independent experiments. Bars show the average ± SD. (E) Quantification of the number of EdU foci found in mitobulbs relative to those found outside of mitobulbs in the cells quantified in (D). (F) qPCR quantification of mtDNA levels in controls cells and two patient fibroblast lines (P1, P2). Each point represents one independent experiment. Bars show the average ± SD. (G) Quantification of EdU foci in control and DRP1 mutants cells labelled with EdU for 4 hours. Each point represents one cell, with at least 44 cells quantified in 3 independent experiments. Bars show the average ± SD. *** p<0.001 two-sided t-test.

### Mutation in DRP1 affects ER-mitochondria contact sites

Previous studies have shown that mtDNA replication initiation occurs at sites where mitochondria are in contact with ER tubules and that loss of ER tubules impairs mtDNA replication ^18,19^. To determine whether fission defects alter mitochondria-ER interaction, we measured ER-mitochondria contact sites in DRP1 mutant primary fibroblasts. We performed a proximity ligation assay (PLA) for Calnexin (general ER marker), and TOM20 (mitochondria) to identify mitochondria-ER contact sites. To assess the specificity of the interaction, we also co-labeled the cells for Calnexin and TOM20. Clear PLA foci were present at sites where Calnexin and TOM20 signals overlapped (Figure 2A; IgG control and Full images in Figure S1B), indicating the identification of authentic ER-mitochondria contact sites. We observe a significant increase in PLA foci in DRP1 mutants (Figure 2B, Full images in Figure S1B), suggesting an increased ER-mitochondria contact sites. A similar increase in PLA foci was also observed when using the well characterized IP3R-VDAC PLA pair (Figure 2C).

**Figure 2.**
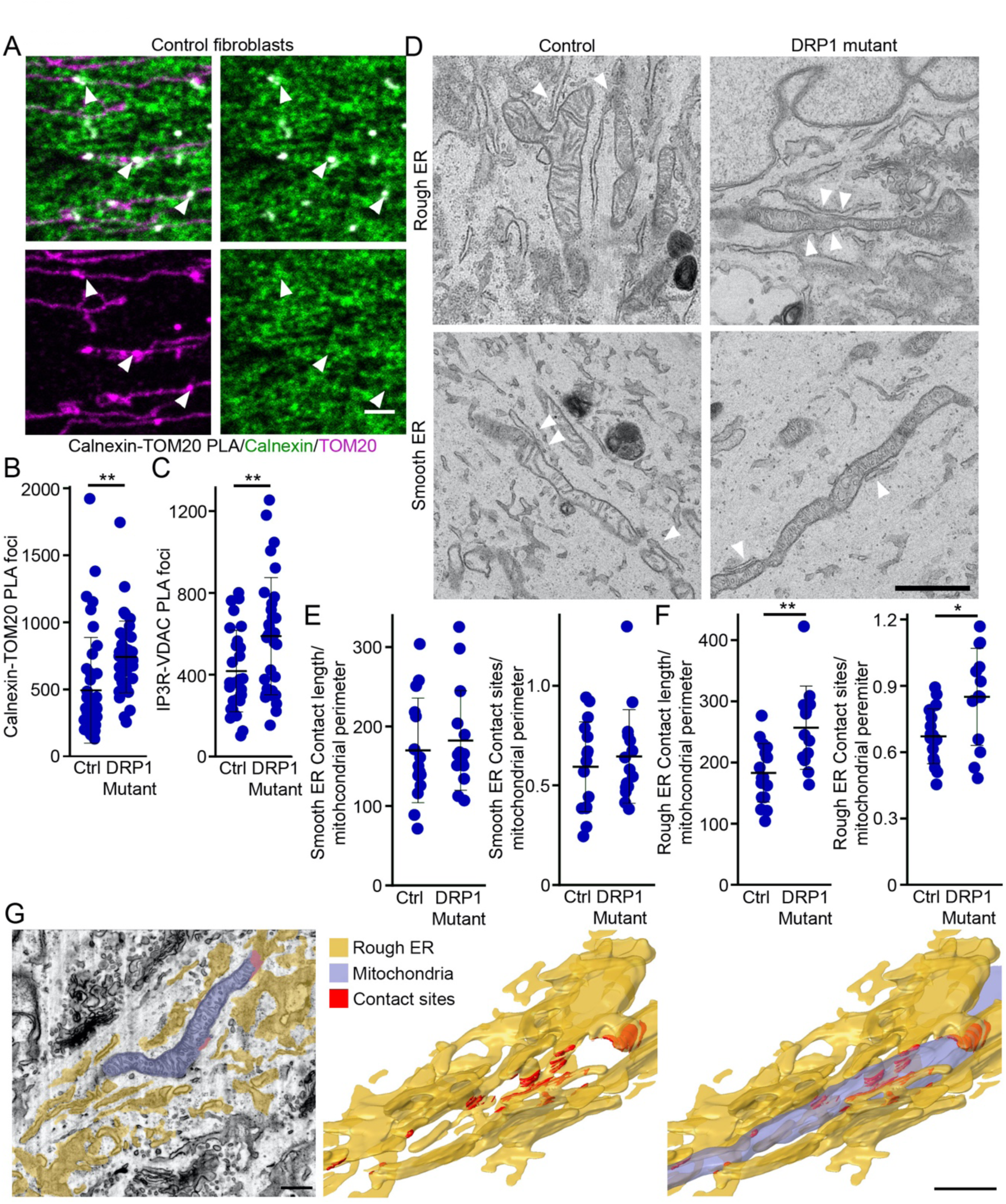
Increased contact sites between rough ER and mitochondria in DRP1 mutant fibroblasts. (A) Representative images of control fibroblasts showing the PLA for Calnexin and TOM20 (white), along with Calnexin (ER, green), and TOM20 (mitochondria, magenta). Arrowheads denote PLA foci. Scale bar 2 µm. Full image in Figure S1 (B) Quantification of Calnexin-TOM20 PLA. Each data point represents one cell. Bars represent the average of 40 cells per genotype in 3 independent experiments ± SD ** p<0.01 two-sided t-test (C) Quantification of IP3R-VDAC PLA. Each data point represents one cell. Bars represent the average of 30 cells per genotype in 3 independent experiments ± SD ** p<0.01 two-sided t-test (D) Representative TEM images of control and DRP1 mutant fibroblasts. Arrowheads denote ER-mitochondria contact sites. Scale bar 1 µm. (E) Quantification of the ERMCs length (µm) per mitochondrial perimeter (Left) and number of ERMCS per mitochondrial perimeter (Right) in TEM images of smooth ER. Each data point represents one cell. Bars represent the average of 15 cells per genotype ± SD. (F) Quantification of the ERMCs length (µm) per mitochondrial perimeter (Left) and number of ERMCS per mitochondrial perimeter (Right) in TEM images of rough ER. Each data point represents one cell. Bars represent the average of 15 cells per genotype ± SD * p<0.05, ** p<0.01 two-sided t-test. (G) FIB-SEM images of a DRP1 mutant mitochondria showing its association with rough ER. Left, FIB-SEM image, middle and right, 3D reconstruction. Scale bar 500 nm.

To further confirm changes in ER-mitochondrial interaction, we measured ER-mitochondria contact sites using transmission electron microscopy (TEM). As mitochondrial fission has been associated with ER tubules ^17–19^ which are generally correlated with smooth ER as found by TEM ^32–35^, we first quantified the interaction between mitochondria and smooth ER. However, there was no difference in smooth ER-mitochondria contact sites between control and DRP1 mutant fibroblasts (Figure 2D-E). While previous studies have mostly focused on the interaction between mitochondria and smooth ER/ER tubules, mitochondria can also interact with rough ER^29,30,36^.

This interaction can in fact readily be observed in control fibroblasts (Figure 2D). Importantly, this interaction was further enhanced in DRP1 mutant fibroblasts, as shown by the increase in both the number of ERMCS per mitochondrial length and the length of these ERMCS (Figure 2D, F). Extensive interactions between mitochondria and rough ER were also evident in a FIB-SEM analysis of DRP1 mutant fibroblasts (Figure 2G). Altogether, our data indicates that ER-mitochondria contact sites are increased in DRP1 mutants and that this increase is mainly contributed by rough ER-mitochondria contact sites.

### A global increase in ER sheets-mitochondrial interaction in DRP1 mutants

Rough ER, as identified by TEM, mostly correlates with ER structures termed ER sheets ^32–35^, which is also supported by our FIB-SEM analysis of rough ER-mitochondria contact sites (Figure 2G). As ER sheets are mostly enriched in the perinuclear area where mitobulbs are also found, we measured the interaction between mitochondria and ER sheets using PLA. For this, we used the ER sheet marker CLIMP63 and the mitochondrial marker TOM20. As shown in Figure 3A-B (Full cells in Figure S2A), DRP1 mutants had significantly increased mitochondria-ER sheets interaction compared to control cells. To further validate the change we observed in ER sheet-mitochondria interaction, we used super-resolution structured illumination microscopy (SIM) of cells labeled with mitotracker orange (mitochondria) and CLIMP63 (ER sheets) (Figure 3C, Full cells in Figure S2B). Consistent with the PLA results, there was an increased colocalization between CLIMP63-labeled ER sheets and mitochondria in DRP1 mutant cells, as quantified using Manders’ coefficient (Figure 3D). The specificity of the measure was validated by rotating one of the channels 90° before measuring Manders’ coefficient ^37^(Figure 3D). As Manders’ coefficients can be affected by structural changes in the organelles tested ^38^, we also measured the overlap between the two structures (see methods) and observed a similar increase in interaction (Figure 3E). Altogether, our data demonstrate that the interaction between CLIMP63-positive ER sheets and mitochondria is increased in DRP1 mutants.

**Figure 3.**
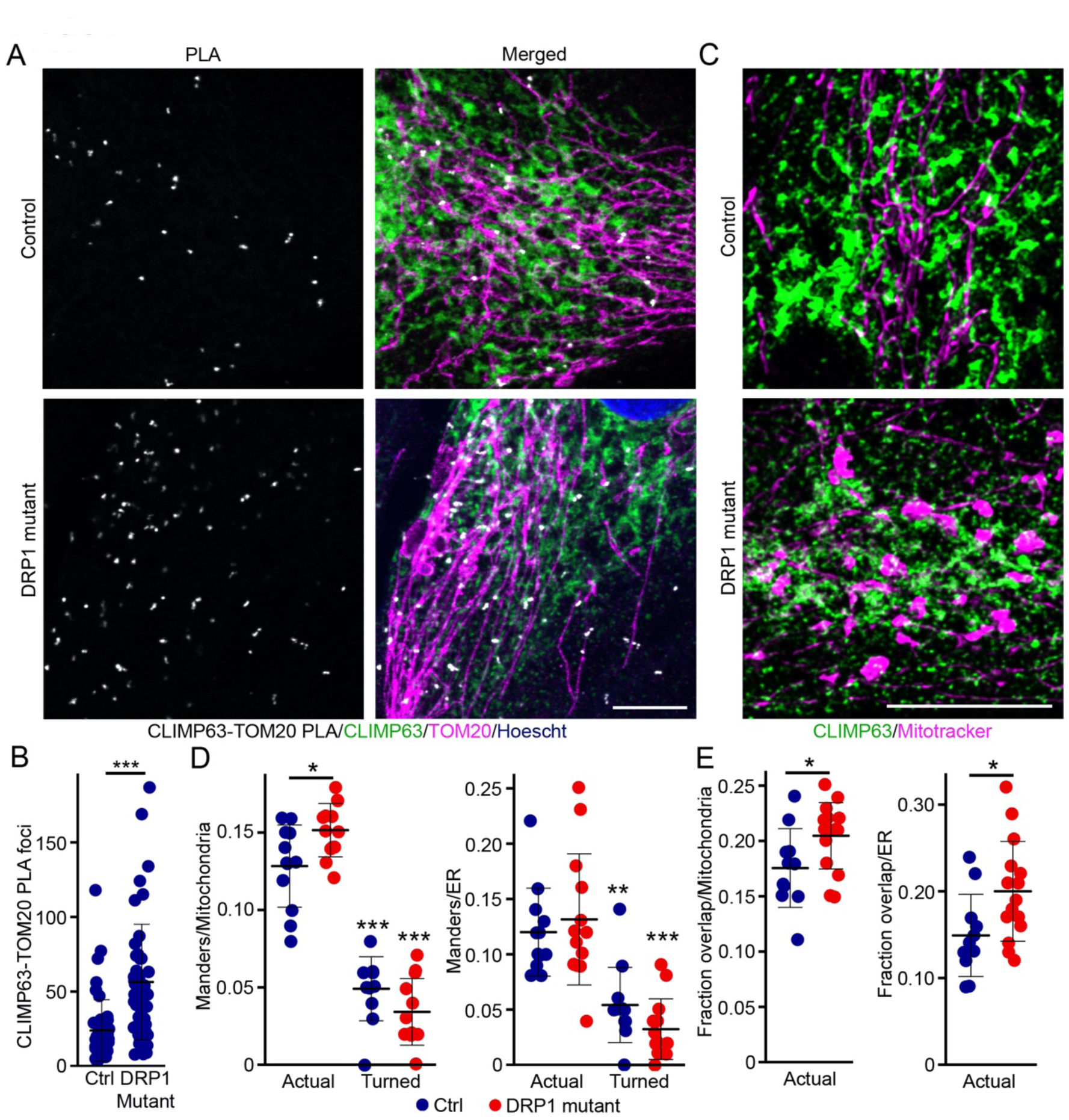
Increased contact sites between CLIMP63-labelled ER and mitochondria in DRP1 mutant fibroblasts. (A) Representative images of control and DRP1 mutant fibroblasts showing the PLA for CLIMP63 and TOM20 (White), along with CLIMP63 (ER sheets, green), TOM20 (mitochondria, magenta) and nuclei (Hoechst, blue). Scale bar 10 µm. Full image in Figure S2A (B) Quantification of CLIMP63-TOM20 PLA. Each data point represents one cell. Bars represent the average of 50 control and 48 mutant cells in 3 independent experiments ± SD *** p<0.001 two-sided t-test. (C) Representative SIM images of control and DRP1 mutant fibroblasts stained for CLIMP63 (ER sheets, green) and Mitotracker orange (mitochondria, magenta). Scale bar 10 µm. Full image in Figure S2B (D) Manders’ coefficients calculated from the SIM images in (C). Left, M1 (relative to mitochondria); Right, M2 (relative to the ER). In the turned condition, the CLIMP63 images were rotated 90° to represent a random distribution. Each data point represents one cell. Bars represent the average of 11 cells per genotype ± SD * p<0.05 One-way ANOVA (E) Fraction of overlapping signal between ER and mitochondria normalized to mitochondria (Left) or the ER area (Right) in SIM images (C). Each data point represents one cell. Bars represent the average of 11 cells per genotype ± SD * p<0.05 two-sided t-test.

To determine if the changes in ER sheet-mitochondria interactions are found in other instances of DRP1 loss of function, we knocked down DRP1 in mouse embryonic fibroblasts (MEFs) (Figure S3A), leading to the formation of a highly elongated mitochondrial network (Figure S3B-C), and measured CLIMP63-TOM20 PLA. Consistent with the loss of DRP1 function altering ER-mitochondria contact sites, the size of individual foci was greater in knocked down cells (Figure S2E), although the number of PLA foci was decreased (Figure S2D). Given the difference in CLIMP63-TOM20 PLA in mutant human fibroblasts and MEFs, we asked if this is the result of different cell contexts or because DRP1 deletion acts differently than the expression of a dominant-negative mutant. To test this, we measured CLIMP63-TOM20 PLA in MEFs expressing a well characterized dominant-negative DRP1 mutant (DRP1K38A) which causes mitochondrial elongation and mitobulb formation (Figure S1A) ^39^. Consistent with the human fibroblast data, DRP1K38A increased the number of CLIMP63-TOM20 PLA foci (Figure S3F). Altogether, our data suggest that DRP1 defect or loss of its function results in alteration in ER sheets-mitochondrial contact sites.

### Differential regulation of ER sheets-mitochondria interaction in DRP1 mutants

ER-mitochondria contact sites are mediated by several protein interaction partners present in the ER and mitochondrial outer membrane. While most of these pairs are expected to be present in both ER tubules and sheets, RRBP1 is an ER sheet-specific protein ^40–42^. To determine if specific ER sheet-mitochondria protein tethers were altered in DRP1 mutant fibroblasts, we performed PLA for RRBP1 and its mitochondrial binding partner SYNJ2BP. In contrast to our TEM and PLA results using general ER (calnexin, IP3R) and ER sheet (CLIMP63) markers, there were fewer RRBP1-SYNJ2BP contact sites in DRP1 mutants (Figure 4A-B, Full image in Figure S4A) compared to control cells. To further confirm this result, we imaged mitochondria (mitotracker orange) and ER sheets (RRBP1) by SIM (Figure 4C, Full image in Figure S4B). Consistent with the PLA data, we observed a significant decrease in the interaction between RRBP1-positive ER sheets and mitochondria as measured by Manders’ coefficients and the overlap between the two organelles (Figure 4D-E). Altogether, our data indicate that while DRP1 mutant fibroblasts show an overall increase in ER sheet-mitochondria contact sites, the specific interaction between the ER sheet resident protein RRBP1 and its mitochondrial partner SYNJ2BP is decreased.

**Figure 4.**
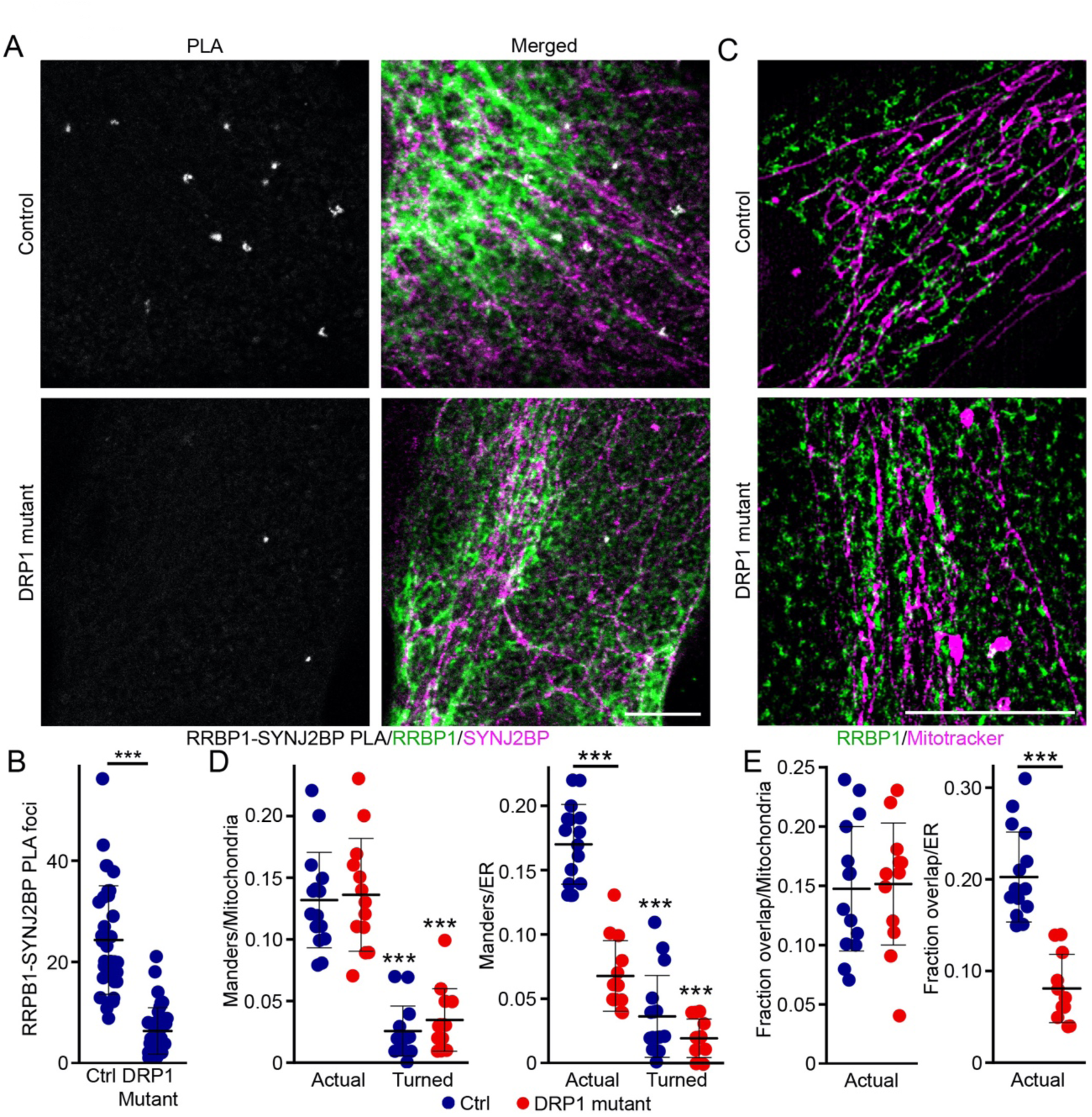
Decreased RRPB1-SYNJ2BP contact sites in DRP1 mutant fibroblasts. (A) Representative images of control and DRP1 mutant fibroblasts showing the PLA for RRBP1 and SYNJ2BP (White), along with RRBP1 (ER sheets, green) and SYNJ2BP (mitochondria, magenta). Scale bar 10 µm. Full image in Figure S4A (B) Quantification of RRBP1-SYNJ2BP PLA. Each data point represents one cell. Bars represent the average of 50 control and 48 mutant cells in 3 independent experiments ± SD *** p<0.001 two-sided t-test. (C) representative SIM images of control and DRP1 mutant fibroblasts stained for RRBP1 (ER sheets, green) and Mitotracker orange (mitochondria, magenta) Scale bar 10 µm. Full image in Figure S4B (D) Manders’ coefficients calculated from the SIM images in (C). Left, M1 (relative to mitochondria); Right, M2 (relative to the ER). In the turned condition, the RRBP1 images were rotated 90° to represent a random distribution. Each data point represents one cell. Bars represent the average of 11 cells per genotype ± SD * p<0.05 One-way ANOVA (E) Fraction of overlapping signal between ER and mitochondria normalized to mitochondria (Left) or the ER area (Right) in SIM images (C). Each data point represents one cell. Bars represent the average of 11 cells per genotype ± SD * p<0.05 two-sided t-test.

### ER sheets are associated with mitobulbs in DRP1 mutants

As ER sheets are predominantly present in the perinuclear region of the cell where mitobulbs are also present, we next determined if ER sheets are associated with mitobulbs. To test this, we labeled control and DRP1 mutant fibroblasts for mitochondria (Mitotracker orange), ER sheets (CLIMP63), and ER (RTN4) and imaged them by confocal microscopy (Figure S5). Mitobulbs were present in the perinuclear region associated with ER sheets, but almost completely absent from the peripheral region characterized by RTN4-positive ER tubules (Figure S5). In fact, the vast majority of mitobulbs marked with the ATP synthase subunit ATP5A were closely associated with ER sheets (Figure 5A, quantification in B), supporting the idea that ER sheets interact with mitobulbs. To further confirm the presence of physical interactions between mitobulbs and ER sheets, we performed PLA for CLIMP63 (ER sheets) and TOM20 (mitochondria). PLA foci were visible at sites where ER sheets were in close contact with mitobulbs (Figure 5C, quantification in 5D), consistent with an interaction between the two structures. Similar results were observed when Calnexin was used instead of CLIMP63 for the PLA (Figure 5D). Consistent with this, a close association of mitobulbs with CLIMP63-positive ER sheets was observed in SIM images (Figure 5E). Altogether, these results indicate that mitobulbs are closely associated with ER sheets, suggesting a role for this interaction in mitobulb formation.

**Figure 5.**
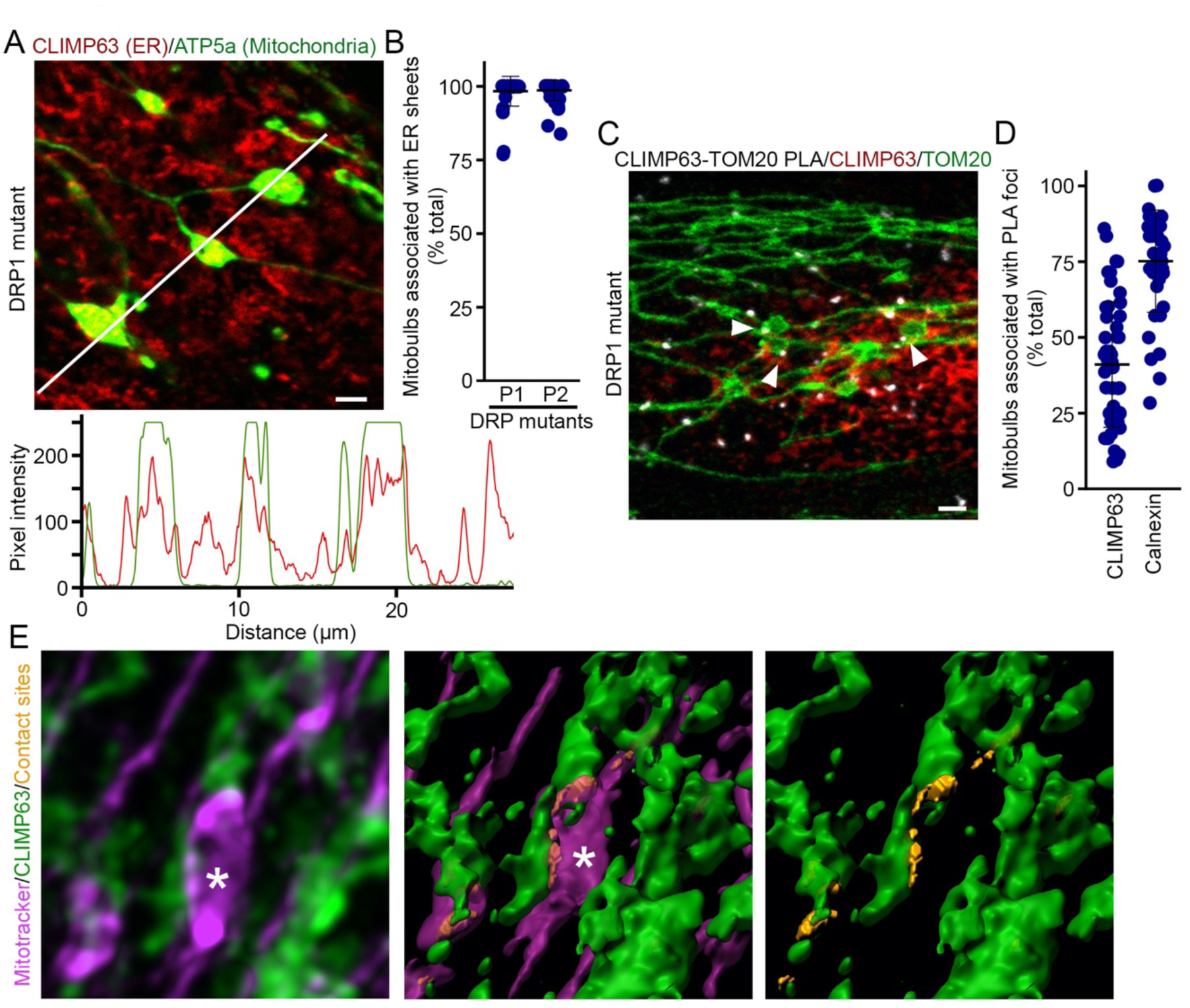
ER sheets are associated with mitobulbs. (A-B) Colocalisation between ER sheets and mitobulbs. Colocalisation was measured in cells immunolabelled for CLIMP63 (ER sheets, red) and ATP5a (mitochondria, green). (A) representative image (Top) and Line scan analysis (Bottom) along the line shown in the image. (B) Quantification of the percent of mitobulbs that are associated with CLIMP63-positive ER sheets in two independent DRP1 mutant lines (P1 and P2). Each data point represents one cell. Bars represent the average of 45 control and 45 mutant mitochondria ± SD. (C-D) Interaction between ER sheets and mitobulbs as measured by PLA for TOM20 and CLIMP63 or Calnexin. (C) Representative image of TOM20-CLIMP63 PLA. Arrowheads denote PLA foci (White) on mitobulbs (TOM20, Green) at sites where they contact ER sheets (CLIMP63, Red). (D) Quantification of mitobulbs associated with PLA foci for TOM20-CLIMP63 and TOM20-Calnexin. Each data point represents one cell. Bars represent the average of 48 (CLIMP63 PLA) and 40 cells (Calnexin PLA) in 3 independent experiments ± SD. Scale bars 2 µm. (E) Representative image of 3D rendering of a SIM image (middle, right) showing a mitobulb (magenta) in association with ER sheets (green) in a DRP1 mutant cell, original image on the left. Contact sites (golden yellow) as identified by the Imaris software. The asterisk denotes a mitobulb.

### Mutation or knockdown of DRP1 alters ER sheet structure

As our results indicate that the interaction between mitochondria and ER sheets is altered in DRP1 mutant fibroblasts, we determined whether this could be associated with alterations in ER sheet structure. For this, we immunolabelled control and DRP1 mutant cells for the ER sheets marker CLIMP63 and found that DRP1 mutants had altered ER sheets compared to control cells. These alterations were characterized by a punctate appearance mostly apparent towards the periphery, and sometimes thick patches of sheets in the perinuclear region (Figure 6A, an enlarged image showing ER sheet structure; complete cell in Figure S5). The overall area covered by ER sheets was also expanded in DRP1 mutant fibroblasts (Figure 6B). Similar alterations were observed in ER sheets labelled with a second ER sheet marker, RRBP1 (Figure 6C). Overall, 80% of mutant cells showed an altered ER sheet phenotype (Figure 6D). CLIMP63 and RRBP1-positive ER sheets also had a fragmented and more punctate appearance when imaged by SIM (Figure 6E, quantification in F).

**Figure 6.**
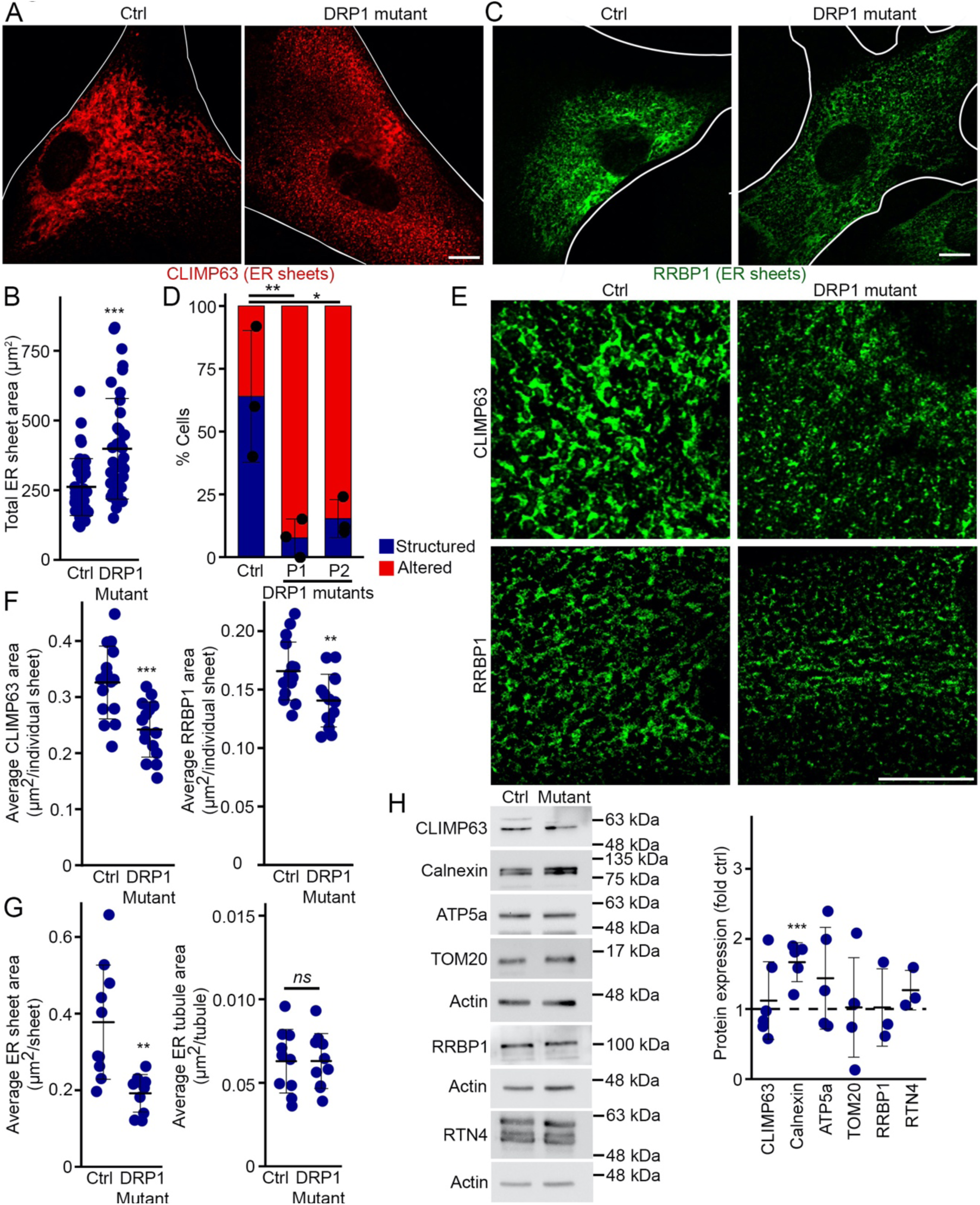
Altered ER sheet structure in DRP1 mutants. (A) Representative images of CLIMP63 staining in control and DRP1 mutant fibroblasts. The white lines denote the edge of the cell as determined by DIC. Full image in Figure S5. Scale bars 10 µm. (B) Quantification of total area covered by ER sheets. Each data point represents one cell. Bars represent the average of 41 cells in 3 independent experiments ± SD *** p<0.001 two-sided t-test. (C) Representative images of RRBP1 staining in control and DRP1 mutant fibroblasts. The white lines denote the edge of the cell as determined by DIC. Scale bars 10 µm. (D) Quantification of ER sheet structure as Structured (Blue) or Altered (red; presence of punctate structures and thick ER sheet patches). Each point represents one independent experiment, with at least 20 cells quantified per experiment. Bars show the average ± SD. * p<0.05, ** p<0.01 One-way ANOVA using the data for Structured ER. (E) Representative SIM images of CLIMP63 (Top) or RRBP1 (Bottom)-labelled ER sheets in Control and DRP1 mutant fibroblasts. (F) Quantification of the average size of individual CLIMP63-labeled (right), RRBP1-labeled (left) ER sheets from SIM images in (E). Each data point represents one cell. Bars represent the average of 15 cells in 3 independent experiments ± SD *** p<0.001 two-sided t-test. (G) Quantification of the average ER sheet (Left) and ER tubule (Right) area in TEM images (Figure 2D). Each data point represents one cell. Bars represent the average of 10 cells ± SD ** p<0.01 two-sided t-test. (H) WB showing ER (CLIMP63, RTN4, RRBP1, calnexin) and mitochondrial proteins (TOM20, ATP5a) in control and DRP1 mutant human fibroblasts. Quantification is shown in the panel on the right, with data point representing expression level of the indicated proteins in DRP1 mutant cells relative to control cells for each experiment. The dashed line shows expression level in control. Bars represent the average of 3-6 experiments ± SD *** p<0.001 two-sided t-test relative to control cells.

To further confirm the structural alteration in ER sheets, we measured the ER area in our TEM images (Figure 2D), where ER sheets were identified as rough ER densely covered with ribosomes. ER sheets from DRP1 mutants had reduced surface area compared to the control (Figure 6G), supporting our data. On the other hand, ER tubule area (identified as smooth ER, free of ribosomes) were similar in control and DRP1 mutants (Figure 6G), consistent with the absence of alterations in ER-mitochondria contact sites for these structures (Figure 2E). Altogether, our data indicate that the structure of ER sheets is altered in DRP1 mutants. Nevertheless, protein levels of CLIMP63, RRBP1, the general ER marker RTN4, and the mitochondrial proteins TOM20 and ATP5a were similar between control and DRP1 mutant cells (Figure 6H). Only Calnexin levels were somewhat increased in mutant cells (Figure 6H). The changes induced by DRP1 mutation are thus unlikely to be caused by changes in protein expression levels (Figure 6H).

Altogether our results indicate that loss of DRP1 function alters ER sheet structure. As we also observed changes in ER sheet-mitochondria interactions in MEFs where DRP1 was knocked down (Figure S3D-E), we then labeled these cells with CLIMP63 (ER sheets) and assessed ER structure. Like patient fibroblasts, DRP1 KD MEFs had mostly punctate ER sheets (Figure S3B, G-H) that covered an expanded area of the cell (Figure S3H). Thus, the presence of mitobulbs is associated with altered ER sheet structure and mitochondrial contacts.

### ER sheets are required for nucleoid maintenance

As there was a correlation between the presence of mitobulbs and ER sheet alterations, we next determined whether these alterations are responsible for mitobulb formation. For this, we used U2OS cells, a cell line that can easily be manipulated. We first confirmed that DRP1 knockdown in these cells leads to ER sheet alterations (Figure S6) and mitobulb formation (Figure 7A-B). We then tested the role of ER sheets in mitobulb formation by manipulating CLIMP63, which is known to alter ER sheet structure and distribution (Figure S6) ^43,44^. Interestingly, while CLIMP63 knockdown did not alter nucleoid size, it led to a decrease in overall nucleoid numbers (Figure 7C-D), indicating that that proper ER sheet structure is required for nucleoid maintenance. CLIMP63 knockdown did not, however, cause mitobulb formation (Figure 7B), indicating that while ER sheets are required to maintain nucleoid numbers, their alterations are not sufficient to cause nucleoid aggregation and mitobulb formation.

**Figure 7.**
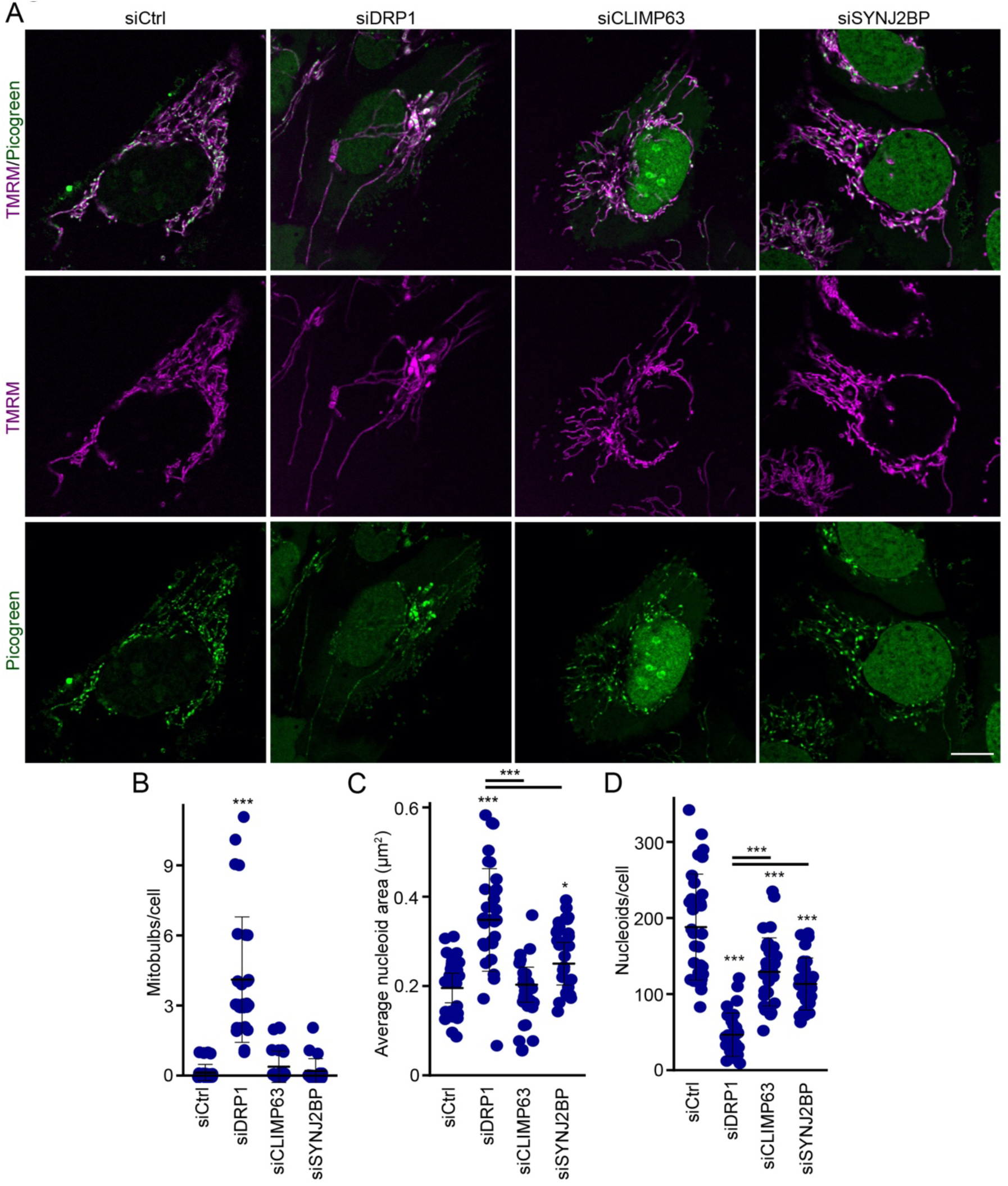
ER sheets are required for proper nucleoid maintenance. (A) Representative images of U2OS cells knocked down for the indicated protein and stained for the Mitotracker (mitochondria, magenta) and Picogreen (nucleoids, Green). Western blots showing altered protein expression in Figure S6 (B) Quantification of the number of mitobulbs present in cells with the indicated protein knocked down by siRNA as in (A). To identify mitobulbs, cells were stained with TOM20 and TFAM. Each data point represents one cell. Bars represent the average of 30 cells in 3 independent experiments ± SD. One way ANOVA. *** p<0.001. (C-D). Quantification of nucleoid size (C) and number (D) in cells as in (A). Each data point represents one cell. Bars represent the average of 30 cells in 3 independent experiments ± SD. One way ANOVA. * p<0.05, *** p<0.001.

To further define the role of ER sheets-mitochondrial contact sites in nucleoid maintenance, we knocked down SYNJ2BP, the mitochondrial binding partner of the ER sheet protein RRBP1. RRBP1 and SYNJ2BP form specific contact sites between ER sheets and mitochondria that are decreased in DRP1 mutant cells (Figure 4). As with CLIMP63 knockdown, loss of SYNJ2BP was not sufficient to cause mitobulb formation (Figure 7A-B). Nevertheless, it caused both a decrease in nucleoid numbers and the enlargement of these nucleoids (Figure 7C-D), suggesting that SYNJ2BP is required for proper nucleoid distribution. Altogether, these results indicate that both the maintenance of proper ER sheet structure and specific contact sites with mitochondria are required for nucleoid maintenance. However, as these alterations remain mild compared to the loss of DRP1, mitobulb formation and drastic nucleoid alterations do require the additional disruption of mitochondrial structure by DRP1.

### Modulation of ER sheets recovers the nucleoid defects present in DRP1 mutants

As our results indicate that ER sheets are required for proper nucleoid maintenance, we then asked whether modulating ER sheet structure can rescue the nucleoid defects present in DRP1 mutant fibroblasts. For this, we modulated ER sheet structure by expressing mCherry-tagged CLIMP63 under conditions that did not overtly alter RTN4-positive ER tubules in control cells (Figure S7A) to avoid nucleoid defects caused by the loss of ER tubules ^19^. Importantly, CLIMP63 expression did not rescue the fused mitochondrial phenotype of the DRP1 mutant (Figure S7B). Nevertheless, CLIMP63 expression altered ER sheet structure resulting in an expanded web-like ER sheet network in both control and DRP1 mutant fibroblasts (Figure S8A-B). Importantly, this was accompanied by a loss of the punctate structures typically observed in mutant cells transfected with mCherry alone (Figure S8B), suggesting that CLIMP63 expression can at least partially rescue the ER sheet defects observed in DRP1 mutants. To determine if this was associated with a change in ER sheets-mitochondria interactions, we performed PLA for CLIMP63 (ER sheets) and TOM20 (mitochondria). CLIMP63 expression did not affect mitochondria-ER sheet interaction in control cells (Figure 8A) but rescued the excessive contact sites found in DRP1 mutant cells (Figure 8A). Similarly, the number of mitobulbs associated with PLA foci was reduced in DRP1 mutant cells expressing CLIMP63 (Figure 8B), indicating that recovery of ER sheet structure in DRP1 mutants normalizes its interaction with mitochondria.

**Figure 8.**
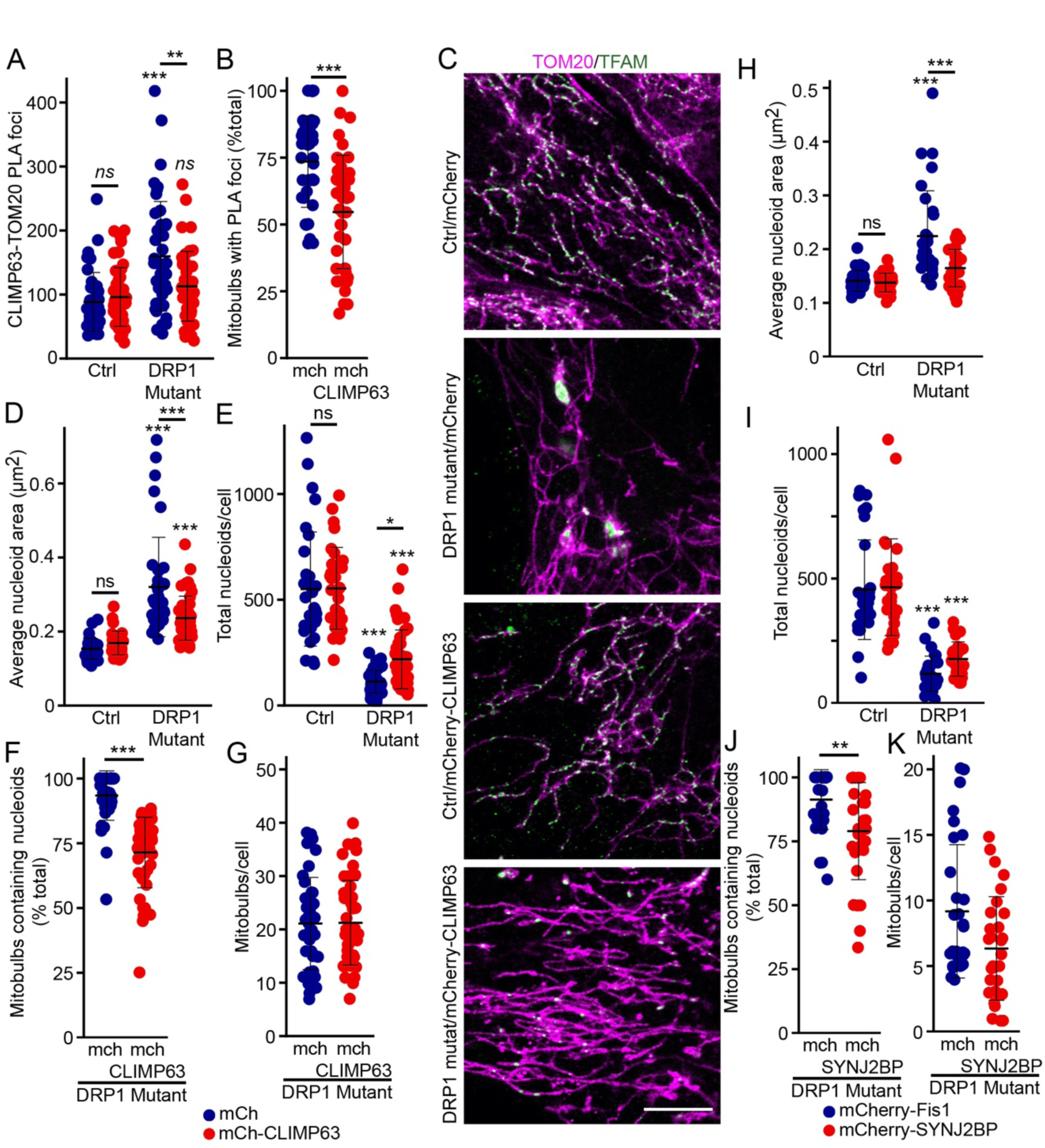
Modulation of ER sheets-mitochondria contact sites rescues nucleoid numbers in DRP1 mutant fibroblasts. (A-B) Quantification of PLA foci (CLIMP63-TOM20) in mCherry and mCherrry-CLIMP63 expressing control and DRP1 mutant cells. Total PLA foci (A) and mitobulbs associated with PLA foci (B) were quantified. Each point represents one cell, with at least 43 cells quantified per condition in 3 independent experiments. Bars show the average ± SD. ** p<0.01, *** p<0.001, ns not significant. One-way ANOVA (C) Representative images of mCherry and mCherrry-CLIMP63 expressing control and DRP1 mutant cells stained for the mitochondrial marker TOM20 (mitochondria, magenta) and the nucleoid marker TFAM (nucleoids, Green). Full images in Figure S9A. Scale bar 10 µm. (D) Quantification of nucleoid area in mCherry and mCherry-CLIMP63 expressing control and DRP1 mutants from images in Figure S9A. Each point represents one cell, with at least 32 cells quantified per condition in 3 independent experiments. Bars show the average ± SD. *** p<0.001, ns not significant. One-way ANOVA. (E-G) Rescue of nucleoid numbers in CLIMP63-expressing DRP1 mutant cells. Quantification of total nucleoids (TFAM-positive, E), mitobulbs containing nucleoids (TFAM-positive, F) and mitochondrial bulb-like structures (independently of the presence of nucleoids, G) in mCherry and mCherrry-CLIMP63 expressing control and DRP1 mutant cells. Each point represents one cell, with 40 mCherry (mch) and 47 mCherry-CLIMP63 cells quantified in 3 independent experiments. Bars show the average ± SD. One way ANOVA (E), two-sided t-test (F, G). * p<0.05, *** p<0.001, ns, not significant. (H) Quantification of nucleoid area in mCherry-Fis1 and mCherry-SYNJ2BP expressing control and DRP1 mutants from images in Figure S9B. Each point represents one cell, with at least 30 cells quantified per condition in 3 independent experiments. Bars show the average ± SD. *** p<0.001, ns not significant. One-way ANOVA. (I-K) Nucleoid numbers in mCherry-SYNJ2BP-expressing DRP1 mutant cells. Quantification of total nucleoids (TFAM-positive, I), mitobulbs containing nucleoids (TFAM-positive, J) and mitochondrial bulb-like structures (independently of the presence of nucleoids, J) in mCherry-Fis1 and mCherrry-SYNJ2BP expressing control and DRP1 mutant cells. Each point represents one cell, with 30 cells quantified in 3 independent experiments. Bars show the average ± SD. One way ANOVA (I), two-sided t-test (J, K). ** p<0.01, *** p<0.001.

Our data suggest that loss of DRP1 results in enlarged nucleoids (Figure 1A) that are likely caused by nucleoid aggregation ^14–16^ as evident from multiple EdU foci in mitobulbs (Figure 1D). To determine the effect of modulating ER sheets on this nucleoid aggregation, we transiently transfected control and DRP1 mutant fibroblasts with mCherry or mCherry-CLIMP63 and immunolabelled them for mitochondria (TOM20) and nucleoids (TFAM). We first determined the effect of the mCherry-CLIMP63 expression on nucleoid aggregation by measuring nucleoid size. Nucleoid size was significantly reduced in DRP1 mutants expressing mCherry-CLIMP63 compared to cells expressing only mCherry (Figure 8C-D, Full cell in Figure S9A), consistent with CLIMP63 expression stimulating nucleoid distribution out of mitobulbs. We then reasoned that if CLIMP63 rescues nucleoid distribution, it should also rescue the decreased nucleoid numbers present in DRP1 mutant cells ^14^. Indeed, while mCherry-CLIMP63 expression did not significantly affect nucleoid numbers in control cells, it caused a significant increase in overall nucleoid content in DRP1 mutants (Figure 8E).

The enlarged nucleoids present in DRP1 mutant cells are found within mitobulbs, we thus then determined the effect of the mCherry-CLIMP63 expression on the presence of mitobulbs. While almost all mitobulbs present in mCherry-transfected DRP1 mutant fibroblasts contained TFAM-positive nucleoids, mCherry-CLIMP63 expression drastically reduced this number (Figure 8F). On the other hand, the total number of enlarged, bulb-like mitochondrial structures was unchanged (Figure 8G), suggesting that DRP1 is still required to maintain mitochondrial structure in these conditions. Altogether, these results indicate that modulating ER sheet structure rescues nucleoid aggregation in DRP1 mutants.

As SYNJ2BP-RRBP1 contact sites are decreased in DRP1 mutant cells (Figure 4) and loss of SYNJ2BP expression impairs proper nucleoid maintenance (Figure 7), we then tested whether increased SYNJ2BP expression could rescue the nucleoid defects present in DRP1 mutant cells. To do this, we expressed either mCherry targeted to mitochondria (mCherry-Fis1) or mCherry-SYNJ2BP which also localizes to mitochondria (Figure S9B). Similar to CLIMP63, expression of mCherry-SYNJ2BP partially rescued nucleoid defects in DRP1 mutant cells but did not alter nucleoid number or size in control cells (Figure 8H-I). Overall, our results indicate that ER sheets and proteins required for ER sheet-mitochondria contact sites regulate nucleoid maintenance.

As SYNJ2BP expression partially rescued nucleoids in DRP1 mutant cells, we also determined whether it promoted nucleoid redistribution similar to CLIMP63. mCherry-SYNJ2BP-expressing DRP1 mutant cells did decrease the number of nucleoid-containing bulb-like structures, but not to the extent observed in mCherry-CLIMP63 expressing cells (Figure 8F, J). This is because the decrease in nucleoid-containing mitobulbs was paralleled by a similar (although not statistically significant) change in total mitobulbs (Figure 8K), suggesting that SYNJ2BP could play a role in mitobulb formation, along with the essential role of DRP1 in this process. Altogether, our data indicate that nucleoid segregation is regulated by ER sheet-mitochondria interaction independently of DRP1-dependent fission.

### Altering ER sheets in DRP1 mutant promotes mtDNA replication and distribution

While our results are consistent with CLIMP63 expression promoting nucleoid distribution away from mitobulbs, it remained possible that it also rescued mtDNA replication defects in DRP1 mutants (Figure 1G). Thus, we measured the effect of CLIMP63 expression on mtDNA replication. Replicating DNA was labeled with EdU, after which cells were directly fixed, immunolabelled for mitochondria (TOM20), and imaged by confocal microscopy. In cells expressing mCherry, DRP1 mutants had fewer EdU-positive nucleoids compared to control cells (Figure 9A-B, Full image in Figure S10), suggesting that DRP1 is required for proper mtDNA replication. Importantly, the expression of CLIMP63 rescued the number of EdU foci in DRP1 mutant cells (Figure 9A-B), suggesting that the effect of DRP1 on nucleoid replication is the consequence of altered ER sheet-mitochondria interaction. On the other hand, CLIMP63 expression did not significantly alter the number of EdU foci in control cells (Figure 9B), indicating that modulating ER sheet structure does not directly affect mtDNA replication.

**Figure 9.**
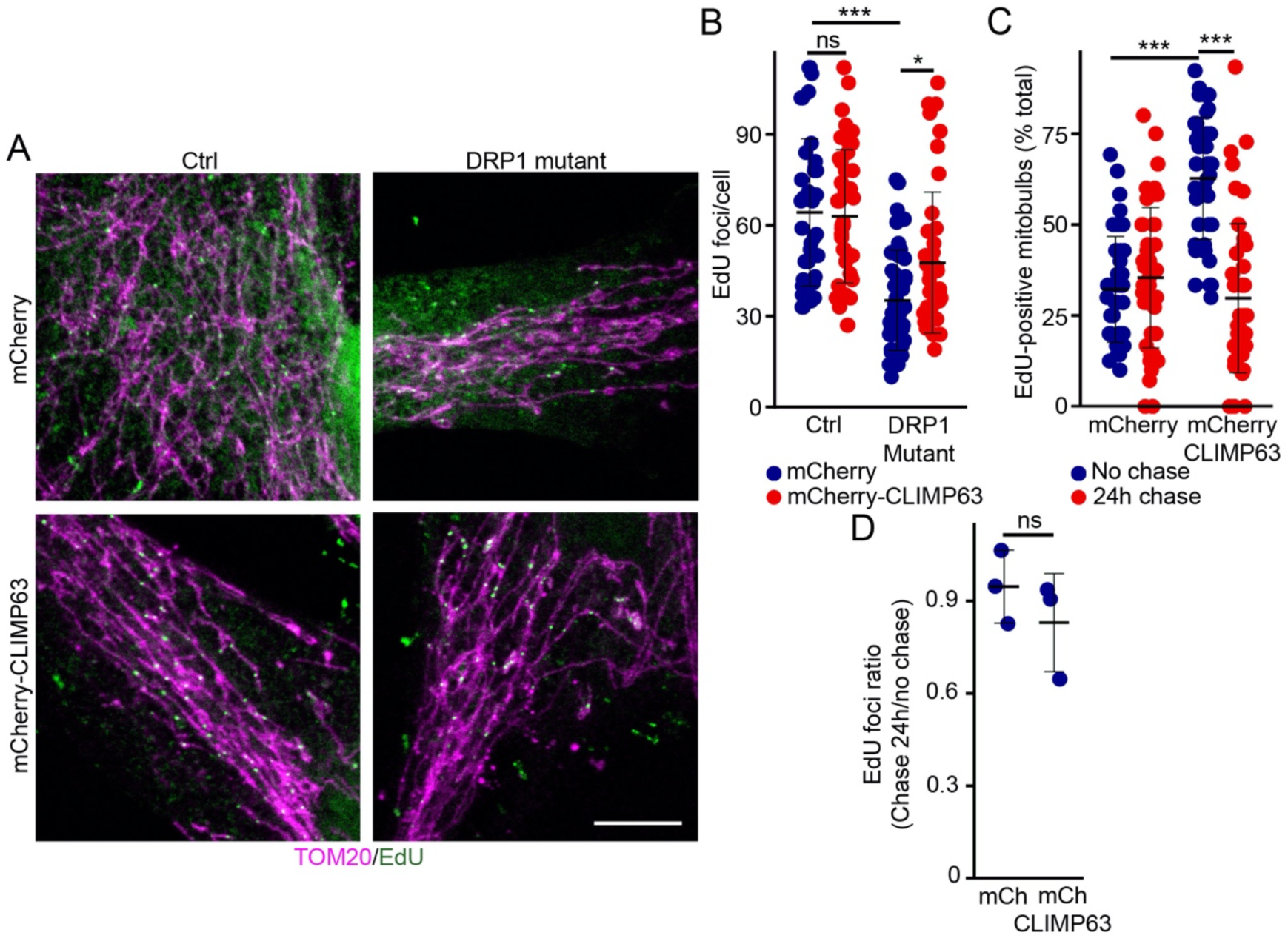
CLIMP63 expression rescues nucleoid replication in DRP1 mutant fibroblasts. (A) Representative images of mCherry and mCherrry-CLIMP63 expressing control and DRP1 mutant cells stained for EdU (Green) and the mitochondrial marker TOM20 (mitochondria, magenta). Full images in Figure S10. Scale bar 10 µm. (B) Quantification of EdU foci in control and DRP1 mutants expressing mCherry or mCherry-CLIMP63 in cells labelled with EdU for 4 hours. Each point represents one cell, with at least 43 cells quantified in 3 independent experiments. Bars show the average ± SD. One way ANOVA. * p<0.05, *** p<0.001, ns not significant. (C) Quantification of EdU-positive mitobulbs in EdU-labelled DRP1 mutants expressing mCherry or mCherry-CLIMP63. Cells were pulsed with EdU as in (A) then the EdU was chased for 24 hours where indicated. Each point represents one cell, with at least 44 cells quantified in 3 independent experiments. Bars show the average ± SD. One way ANOVA. *** p<0.001. (D) EdU foci ratio (chase/no chase) from the experiments in (C). Each point represents an independent experiment (n=3). Bars show the average ± SD. Two-sided t-test. ns not significant.

As clusters of EdU-positive mtDNA are found within mitobulbs (Figure 1C-D), we then specifically addressed the effect of CLIMP63 expression on mtDNA replication within mitobulbs. Consistent with the total EdU counts, expression of CLIMP63 in DRP1 mutant fibroblasts caused a large increase in EdU incorporation in mtDNA present within mitobulbs (Figure 9C). We then took advantage of the accumulation of EdU foci within mitobulbs to determine whether the reduction in nucleoid size that we observed in CLIMP63-transfected DRP1 mutant cells (Figure 8D) is due to the rescue of nucleoid segregation from mitobulbs towards the rest of the mitochondrial network. To do that, we chased EdU for 24 hours before fixing the cells. Consistent with our observation that the number of nucleoid-containing mitobulbs was decreased in CLIMP63-expressing cells (Figure 8F), the number of EdU-positive mitobulbs was decreased in CLIMP63-expressing DRP1 mutant cells, but not mCherry-expressing cells at 24 hours (Figure 9C). Importantly, this was not due to the loss of overall EdU staining in mutant cells (Figure 9D). Overall, our data indicates that modulating ER sheets-mitochondrial interaction through altering ER sheet structure reactivates both mtDNA replication and nucleoid distribution in DRP1 mutant cells even in the absence of mitochondrial fission. Thus, proper ER sheets-mitochondria interaction is essential to modulate mtDNA replication and nucleoid segregation.

## Discussion

mtDNA replication is associated with mitochondrial dynamics, especially mitochondrial fusion which is required for mtDNA replication and integrity ^11,12^. On the other hand, defects in mitochondrial fission impair nucleoid segregation and result in nucleoid aggregation within mitobulbs ^13,15,16^. With the help of Mitomate Tracker, an automated tool developed in our lab, we have shown that this alters overall nucleoid distribution within mitochondrial networks ^14^. Nevertheless, the mechanism by which nucleoid distribution is altered in cells with fission defects remains unclear. Here, we demonstrate that ER sheet-mitochondria contact sites regulate mtDNA replication and distribution within mitochondrial networks.

ER tubule-mitochondria contact sites play an important role in regulating mitochondrial dynamics and function ^17,18,41^ including mtDNA replication ^19^. Our data shows that ER sheets also regulate mtDNA replication and segregation by directly interacting with mitochondria. This is consistent with the previous observation that ER sheets interact with mitochondria and regulate lipid metabolism in mouse liver cells ^29^, suggesting an important physiological role for this interaction. As we previously showed that, in DRP1 mutants, nucleoids aggregate around the perinuclear region where ER sheets are found ^14^, we further defined how ER sheets-mitochondria interaction impacts this process. Our data demonstrates that cells with defective DRP1 have altered ER sheet structure. Furthermore, DRP1 mutant cells showed increased interaction between ER sheets and mitochondria, including with mitobulbs. Of note, we did not observe distinct changes in ER tubules in our cell models, possibly because ER tubular formation requires the variable domain of DRP1 ^45^ which is still intact in our mutant lines (mutation in the middle domain (G362D))^31^. Nevertheless, as the ER sheet alterations present in DRP1 mutants also occurred in cells in which DRP1 was knocked down, they are likely the consequence of the loss of function of DRP1 rather than a specific defect within one of DRP1’s structural domains.

The effects of DRP1 alterations on ER sheet-mitochondria interactions are complex. While dominant-negative DRP1 mutants (DRP1G362D (mutant cells), DRP1K38A) cause an increase in overall ER sheet-mitochondria interactions as measured by PLA for CLIMP63 and TOM20, as well as TEM and SIM, DRP1 knockdown reduces ER sheets-mitochondria contact sites, consistent with the previous observation^46^. While this suggests that dominant-negative forms of DRP1 act differently than its absence, the key consequence of DRP1 loss of function is likely the loss of SYNJ2BP-RRBP1 interaction. In fact, this interaction was lost in DRP1 mutant cells and deleting SYNJ2BP in U2OS cells partially recapitulated the nucleoid defects found in DRP1 mutant human fibroblasts. Furthermore, expression of SYNJ2BP in DRP1 mutant fibroblasts rescued the nucleoid defects present in these cells, further supporting a crucial role for SYNJ2BP in nucleoid maintenance.

One important conclusion from our work is that the altered ER sheet-mitochondria contact sites present in DRP1 mutant cells are associated with the accumulation of mtDNA within mitobulbs. As a consequence, modulating ER sheets by expressing CLIMP63 rescued ER sheet-mitochondria contact sites, nucleoid aggregation, and nucleoid number in DRP1 mutants. Similarly, expression of SYNJ2BP which has been previously shown to increase ER sheets-mitochondria contact sites^40,47^ rescued nucleoid number and size, supporting the idea that ER sheets-mitochondria contact sites regulate nucleoid distribution. In addition, our EdU data demonstrated that proper ER sheet-mitochondria contact sites are necessary to actively regulate mtDNA replication. Altogether, our data show that ER sheet-mitochondria contact sites are essential for both mtDNA replication and distribution. Furthermore, while the alterations in mitochondrial structure directly caused by the lack of fission is likely required for mitobulb formation, ER sheet-mitochondria contact sites regulate mtDNA maintenance independently of DRP1-dependent mitochondrial fission.

While we demonstrated that ER sheets-mitochondria contact sites play an important role in mtDNA replication and distribution, ER tubules are still required. This was shown in a previous study overexpressing CLIMP63 under conditions that caused the complete loss of ER tubules, which significantly reduced mitochondrial DNA replication in cells with wild-type DRP1 ^19^. However, when we expressed CLIMP63 under conditions that do not affect tubular ER structure, mtDNA replication was unaltered in cells with wild-type DRP1. Furthermore, ER tubule structure and interaction with mitochondria were unaffected in cells with defective DRP1. This further suggests that the regulation of mtDNA replication and segregation by ER sheets is independent of ER tubules.

Our study also provides novel insights into the cellular defects that could underlie the clinical phenotype present in DRP1 mutant patients. Reported human DRP1 mutations are mostly found in the middle and GTPase domains of DRP1, leading to peripheral neuropathy, epilepsy, optic atrophy, and encephalopathy in these patients ^31,48^. Here, we have used patient fibroblasts with a dominant negative mutation in the middle domain of DRP1 [NP_036192.2:(p.Gly362Asp], leading to refractory epilepsy^31^. At the cellular level, conditional deletion of DRP1 in mice affects the movement of mitochondria to the nerve terminal, leading to the loss of dopamine neurons ^49^, while its postnatal deletion in CA1 hippocampal neurons results in functionally impaired mitochondria at the nerve terminals ^50^. Based on our key findings here, we speculate that the altered mtDNA replication and nucleoid distribution could severely hamper the movement of functional mitochondria towards the synaptic nerve terminal in the patient neurons and thereby impairing neuronal signaling. Further research work is required to validate this hypothesis in neuronal cells.

Altogether, our results demonstrate the importance of ER sheets-mitochondria contact sites in mtDNA maintenance and nucleoid distribution. Alteration of these contact sites following DRP1 defect leads to the perinuclear accumulation of mtDNA, which could explain the defects observed in these cells in the absence of overt metabolic alterations ^31^.

## Acknowledgements

This work was supported by grants from the Natural Sciences and Engineering Research Council of Canada and the Fondation de l’UQTR to MG and the Canadian Institutes of Health Research and the Canadian Foundation for Innovation to JV. KAN is supported by a scholarship from Qatar University. HSI was supported by a Queen Elizabeth II Diamond Jubilee Scholarship and an FRQ-NT scholarship. Object Research Systems provided access to Dragonfly software.

## Author contribution

HSI performed the cell biology experiments. HSI and MG designed the experiments. HSI, JDG, SB and MG analysed the data. AL reconstructed 3D FIB-SEM data with direction from FR. KAN performed lattice SIM microscopy and image reconstructions with direction from JV. MAL provided the clinical samples. HSI and MG wrote the paper. All authors reviewed and discussed the manuscript.

## RESOURCE AVAILABILITY

### Lead contact

Further information and requests for resources and reagents should be directed to and will be fulfilled by the lead contact, Marc Germain

### Materials availability

This study did not generate new unique reagents.

### Data and code availability

- All data reported in this paper will be shared by the lead contact upon request.
- This paper does not report original code.
- Any additional information required to reanalyze the data reported in this paper is available from the lead contact upon request.

## EXPERIMENTAL MODEL AND STUDY PARTICIPANT DETAILS

### Primary human fibroblasts

Primary human fibroblasts (controls and DRP1 mutants) were generated from skin biopsies, collected as part of an approved research protocol (Research Ethics Board of the Children’s Hospital of Eastern Ontario (DRP1 mutants)), and written informed consent from participants was obtained. This study was performed in accordance with the Declaration of Helsinki. Biopsy samples were processed as described and cultured in Dulbecco’s modified Eagle’s medium (DMEM) containing 10% fetal bovine serum (FBS), supplemented with Penicillin/Streptomycin (100 IU/ml/100µL/mL)^31^.

### Cell lines

Immortalized Mouse Embryonic Fibroblasts (MEFs)^51^ and U2OS were cultured in DMEM supplemented with 10% fetal bovine serum.

## METHOD DETAILS

### Live cell imaging

For live cell imaging, cells were seeded onto glass bottom dishes and allowed to adhere to the plate. Then cells were stained for 30 minutes with 250 nM TMRM (Thermo fisher Scientific, T668) and the DNA dye PicoGreen (Thermo Fisher Scientific, P11495) (3 µL/mL). After staining, cells were washed 3 times with pre-warmed 1X phosphate buffered saline (PBS), and normal growth media was added prior to imaging.

### siRNA treatment

MEFs were seeded onto 24 well dish and transfected with 15nM of DRP1 siRNA (Thermo Fisher Scientific, Silencer Select, 4390771), and negative siRNA (Thermo Fisher Scientific, Silencer Select, 4390843) using siLenFect lipid reagent (Bio-Rad,1703361). U2OS cells were similarly transfected with 15nM of DRP1 siRNA (Horizondiscovery, siGenome SMART pool siRNA, M-012092-01-0005), CLIMP siRNA (Horizondiscovery, siGenome SMART pool siRNA, M-012755-02-0005), SYNJ2BP siRNA (Horizondiscovery, siGenome SMART pool siRNA, M-021176-01-0005) and negative siRNA using silenfect reagent for 96hrs. After 48 hrs, cells were treated again with siRNA for another 48 hours. The cells were collected for either western blotting or seeded onto the coverslips for immunofluorescence.

### Transient transfections

Primary cells were trypsinized and centrifuged at 577 X g for 10 minutes. Cell pellets were suspended in 10µl of the Neon transfection buffer R (ThermoFisher Scientific, MPK1096). Cells were transiently transfected with mCherry (gift from Michael Davidson, Addgene, 54517), mCherry-tagged CLIMP63 (gift from Gia Voeltz ^52^, Addgene, 136293),mcherry-FIS1 (gift from Uri Manor^53^), mch-SYNJ2BP (gift from Marc Tramier^54^)using the Neon transfection system (ThermoFisher Scientific, MPK5000) according to the manufacturer’s protocol. To minimize effects on ER tubules, minimal concentrations (1µg) of mCherry and mCherry-CLIMP63 plasmids were used. MEF cells were transiently transfected with DRP1 K38A (gift from Alexander van der Bliek & Richard Youle^55^, Addgene, 45161) using Metafectene Pro (Biontex, T040) for 24 hours.

### Immunofluorescence

Cells were seeded onto glass coverslips (Fisherbrand, 1254580) and allowed to adhere overnight. Mitochondria was stained using 50 nM Mitotracker Orange (Thermo fisher scientific, M7510) prior to fixing for certain experiment. Cells were fixed with 4% paraformaldehyde for 15 minutes at room temperature (RT). For antigen retrieval, cells were incubated in sodium citrate buffer (pH 6.0) for 10 minutes at 95°C. Then, cells were permeabilized with 1% BSA / 0.2% Triton X-100 in PBS followed by blocking with 1% BSA / 0.1% Triton X-100 in PBS. The following antibodies were used: CLIMP63 (Rb, Abcam, ab84712, 1:150), CLIMP63 (Mo, ENZO, ENZ-ABS669, 1: 200), RRBP1 (Rb, Novus Biologicals, NBP1-32813, 1:200), TOM20 (Rb, Abcam, ab186735, 1:250), mtTFAM (Mo, Novusbio, NBP1-71648, 1:150), Calnexin (Mo, Millipore, MABF2067, 1:200), RTN4/NOGOA (Rb, Bio-Rad, AHP1799, 1:200), ATP5a (Mo, Abcam, ab14748, 1:150), SYNJ2BP (Rb, Sigma-Aldrich, HPA000866, 1:200), RRBP1 (Mo, Thermo Fisher Scientific, MA5-18302, 1:200), IP3R1 (Rb, Novus Biologicals, NBP2-22458, 1:200), VDAC(Mo, Sigma-Aldrich, MABN504, 1:200). Next, cells were incubated with fluorescent tagged secondary antibody (Jackson Immunoresearch, 1:500).

### Microscopy

Images were acquired with a Leica TSC SP8 confocal microscope fitted with a 63x/1.40 oil objective using the optimal resolution for the wavelength (determined using the Leica software). For super-resolution microscopy, z-stack images were taken using a Zeiss Elyra 7.2 lattice structured illumination microscope (SIM) system fitted with 63X 1.4 na Plan-Apochromat oil objective with 1X (CLIMP63, mitotracker labeled images) or 1.6X optivar (RRBP1, mitotracker labelled images) and dual sCMOS cameras (pco.edge). CLIMP63 and RRBP1 images with a spatial resolution of 90 nm were constructed using the SIM^2^ processing, using an iteration strength appropriate for the signal to noise ratio (3-5) for the CLIMP63 or RRBP1 data. Mitotracker images were processed using standard SIM.

### Image analysis

All image manipulation and analysis were done in Image J/Fiji except where noted. To verify the interaction between organelles (bulb-like mitochondria and ER sheets), line scan analysis was performed using the *RGB profiler* in Image J. To avoid bias during manually classification of ER structure, images were renamed to random numbers and both control and mutant images were shuffled together. The blindfolded ER sheet images were manually classified as structured (properly organized ER sheets) or altered (fragmented and expanded ER sheet structure) using reference images. In experiments where mCherry-CLIMP63 was expressed the category “expanded” was added to denote cells with properly structured sheets that extended towards the periphery of the cell. In addition, we segmented the ER sheet images in ImageJ (Filter/minimum (0.5), Filter/Median (1.0), then thresholding and adjusting the resulting image using Binary/Erode) and measured total area ER sheets using the Analyse Particle function. To measure the area occupied by ER sheets in MEF and U2OS cells, the overall ER sheet and total cell area were measured using ImageJ. Mitochondrial structures were manually quantified by binning them into the indicated categories (short, intermediate, elongated). For the SIM images, the processed images were used to measure ER sheet-mitochondria contact sites using Manders’ colocalization coefficient. Maximum z-projected images were used for the analysis. ER sheets were segmented using Image J *filters* (median (1), mean (25)) and thresholded by default method. Mitochondria were segmented using the Image J tool *Tubeness* and thresholded by default method ^14^. Segmented images were analyzed using the Image J plugin *Just another colocalization plugin (JaCoP)*^56^. The specificity of the measure was validated by rotating one of the channels 90° before measuring Manders’ coefficient^37^. ER sheet-mitochondria contact site area was measured by identifying regions of colocalization in the SIM processed images using the Image J tool *Hue*. The average ER sheet size was measured using segmented ER sheets images using the Image J tool *Analyze particle*. 3D-SIM reconstructions were performed using Bitplane Imaris software v. 9.5.

### Proximity Ligation Assay (PLA)

Cells were grown on glass coverslips and fixed with 4% paraformaldehyde. Following antigen retrieval, cells were permeabilized with 1% BSA/0.2% Triton X-100 in PBS for 15 minutes at RT. The ER was marked with an antibody recognizing CLIMP63 or calnexin or RRBP1 and mitochondria was marked with an antibody recognizing TOM20 or SYNJ2BP. The PLA assays were performed using Duolink In Situ green kit Mouse/ Rabbit (Sigma Aldrich) following the manufacturer’s protocol. The primary antibodies were then labelled using fluorescent-tagged secondary antibodies. Total and mitobulb associated PLA foci were manually counted at the site of CLIMP63-TOM20 and RRBP1-SYNJ2BP signal while overall calnexin-TOM20 and IP3R1-VDAC foci were quantified using the *analyze particles* function in ImageJ.

### Transmission Electron Microscopy (TEM)

Primary fibroblasts were seeded on Nunc Lab-Tek chamber slides (Thermo Fisher Scientific, 177437) and allowed to adhere overnight. Cells were washed with 0.1M sodium cacodylate buffer (pH 7.3) and fixed with 2.5% glutaraldehyde in 0.1M sodium cacodylate buffer (pH 7.3), overnight at 4°C. Fixed cells were further processed in McGill’s facility for electron microscopy research (FEMR). Images were acquired using a EMS208S electron microscope (Philips) by an independent trained operator in University of Quebec in Trois-Rivières’ TEM facility. We manually measured ER surface area for 10 randomly chosen rough ER/smooth ER in each of the four quadrants per image of the cell. Rough ER-mitochondrial contact sites in the perinuclear region were manual quantified, considering only contact sites ≤ 30 nm based on a previous study ^57^. Smooth ER-mitochondria contact sites were manually measured near the cellular periphery per image of the cell.

### Focused Ion Beam Scanning Electron Microscopy

Sample blocks for analysis by FIB-SEM were prepared as for TEM. Each Epon block was trimmed and mounted on a 45° pre-titled SEM stub and coated with a 4-nm thick of Pt to enhance electrical conductivity. Milling of serial sections and imaging of block face after each Z-slice was carried out with the Helios Nanolab 660 DualBeam using Auto Slice & View 4.1 software (FEI Company, Hillsboro, OR USA).

A block was first imaged to determine the orientation relationship between the block face of ion and electron beams. A protective Pt layer 26.3 μm long, 1.5 μm wide and 2 μm thick was deposited on the surface of the region of interest to protect the resin volume from ion beam damaging and correct for stage and/or specimen drift, i.e., perpendicular to the image face of the volume to be milled. Trenches on both sides of the region were created to minimize re-deposition during the automated milling and imaging. Distinct imaging fiducials were generated for both ion beam and electron beam imaging and were used to dynamically correct for any drift in x and y during a run by applying appropriate SEM beam shifts. Milling was performed at 30 kV with an ion beam current of 2.5 nA, stage tilt of −0.5 degree, working distance of 4 mm. With each step, 4 nm thickness of the material was removed with the ion beam. Each newly milled block face was imaged with Through the Lens Detector (TLD) for backscattered electrons at an accelerating voltage of 2 kV, beam current of 0.4 nA, stage tilt of 37.5 degree, and working distance of 2 mm. The pixel resolution was 4.15 nm with a dwell time of 30 μs. Pixel dimensions of the recorded image were 1536 x 1024 pixels. 201 images were collected, and the contrast of the images inversed.

### Processing of FIB-SEM images

Three-dimensional reconstruction of one long mitochondrion and associated ER network was done with Dragonfly software (Ver. 2022.1; Object Research Systems, Montreal QC). Mitochondria and ER from DRP1 mutant patient cells were segmented manually within the painting tool in Dragonfly from 75 slices as 4 nm slice spacing throw 0.3 µm depth. The boolean operation option was used to highlight the intersection regions between mitochondria and ER (contact site regions). The total number of voxels segmented as RE was 8 513 191, 2 337 756 voxels as mitochondria and 56 677 as contact sites between the two organelles.

### Western Blot

Cells were lysed in 10 mM Tris-HCl, pH 7.4, 1mM EDTA, 150 mM NaCl, 1% Triton X-100, 50 mM sodium fluoride, complemented with a protease inhibitor cocktail (Sigma-Aldrich), centrifuged at 15890 X g for 5 minutes and protein supernatants collected. Protein concentrations were estimated calorimetrically by DC protein assay kit (BioRad). For SDS-PAGE, 20 µg (siRNA treatment in MEFs)/25 µg (Mitochondrial and ER proteins in human fibroblasts) of proteins were mixed with 1X Lammeli buffer containing β-mercaptoethanol, then subjected to SDS-PAGE, transferred to nitrocellulose membranes and blotted with the indicated antibodies (DRP1 (BD Transduction Laboratories, 611112, 1:1000), CLIMP63 (Mo, ENZO, ENZ-ABS669, 1: 500), TOM20 (Rb, Abcam, ab186735, 1:1000), Calnexin (Mo, Millipore, MABF2067, 1:1000), RRBP1 (Rb, Novus Biologicals, NBP1-32813, 1:1000), RTN4/NOGOA (Rb, Bio-Rad, AHP1799, 1:1000), ATP5a (Mo, Abcam, ab14748, 1:1000), SYNJ2BP (Rb, Sigma-Aldrich, HPA000866, 1:1000), HRP-tagged Actin (1:10,000). Membranes were then incubated with a 1:5000 dilution of horseradish peroxidase-conjugated secondary antibodies (Jackson Immunoresearch) and visualized by enhanced chemiluminescence (Thermo Fisher scientific) using a Bio-Rad imaging system. Protein expression was measured using Biorad’s Imagelab software and normalized to actin. Data was represented as fold changes relative to control cells.

### mtDNA copy number

DNA was isolated from primary cells using PureLink Genomic DNA mini kit (Thermo Fisher Scientific, K182001). 100 ng of DNA sample was utilized for the quantification of mtDNA copy number using Bio-Rad CFX Real-Time PCR system. Target mtDNA gene (Forward primer: 5′-CACCCAAGAACAGGGTTTGT-3′, Reverse primer: 5′-TGGCCATGGGTATGTTGTTAA-3′, Invitrogen custom primers) and reference 18S ribosomal RNA gene (Forward primer: 5′-TAGAGGGACAAGTGGCGTTC-3′, Reverse primer: 5′-CGCTGAGCCAGTCAGTGT-3′, Invitrogen custom primers) was independently amplified using thermocycling conditions as described in ^58^. Each quantification PCR reaction sample contain template DNA, PowerUp SYBR Green Master Mix (Thermo Fisher Scientific, A25742) and 500 nM of primers (final concentration). The relative mtDNA copy number was assessed using Agilent MxPro – Mx3000P Multiplex Quantitative PCR Systems. Relative mtDNA copy number was quantified by Livak method^59^.

### EdU Labeling

Primary fibroblasts (Control and DRP1 mutants) were incubated with 60µM EdU for 2 hours at 37°C. For the chase experiments, the EdU containing media was replaced with fresh media and incubated further for 24 hours. Cells were then fixed with 4% paraformaldehyde for 15 minutes at RT, permeabilized and EdU detected using Click-iT EdU imaging kit (Thermo Fisher Scientific, C10337). Cells were then immunolabelled for TOM20 (Abcam, ab186735, 1:250). EdU foci in mitobulbs and total EdU foci were manually counted.

## QUANTIFICATION AND STATISTICAL ANALYSIS

All graphs and statistical analysis were done using R. Immunofluorescence data were quantified and images representative of at least three independent experiments shown (exact n are in the quantification figures). Data are represented as average ± SD as specified in figure legends. Statistical significance was determined using Student’s t test (between 2 groups) or one-way ANOVA with a Tukey post hoc test (multiple comparisons).

## KEY RESOURCES TABLE

| REAGENT or RESOURCE | SOURCE | IDENTIFIER |
| --- | --- | --- |
| Antibodies |  |  |
| CLIMP63 | Abcam | ab84712 |
| CLIMP63 | ENZO | ENZ-ABS669 |
| RRBP1 | Novus Biologicals | NBP1-32813 |
| TOM20 | Abcam | ab186735 |
| mtTFAM | Novus Biologicals | NBP1-71648 |
| Calnexin | Millipore | MABF2067 |
| RTN4/NOGOA | Bio-Rad | AHP1799 |
| ATP5a | Abcam | ab14748 |
| SYNJ2BP | Sigma-Aldrich | HPA000866 |
| RRBP1 | Thermo Fisher Scientific | MA5-18302 |
| Fluorescent-tagged secondary antibody | Jackson ImmunoResearch | 711-605-152(Anti-Rabbit 647), 715-605-150 (Anti-mouse 647), 711-165-152(Anti-Rabbit cy3), 711-545-152 (Anti-Rabbit 488), 715-165-150 (Anti-Mouse cy3), 715-545-150 (Anti-Mouse 488) |
| DRP1 | BD Transduction Laboratories | 611112 |
| Horseradish peroxidase-conjugated secondary antibodies | Jackson ImmunoResearch | 115-035-003(Anti-Mouse), 111-035-003(Anti-Rabbit) |
| IP3R1 | Novus Biologicals | NBP2-22458 |
| VDAC | Sigma-Aldrich | MABN504 |
| Chemicals, Peptides, and Recombinant Proteins |  |  |
| Dulbecco's modified Eagle's medium | Wisent | 319-005-CL |
| Trypsin | Wisent | 325-042-CL |
| Penicillin-Streptomycin | Wisent | SV30010 |
| Fetal bovine serum | Corning | 35-077-CV |
| TMRM | Thermo fisher Scientific | T668 |
| PicoGreen | Thermo Fisher Scientific | P11495 |
| Phosphate buffered saline | Wisent | 311-425-CL |
| siLenFect lipid reagent | Bio-Rad | 1703361 |
| Neon transfection buffer R | Thermo Fisher Scientific | MPK1096 |
| Mitotracker Orange | Thermo fisher scientific | M7510 |
| Paraformaldehyde | Sigma-Aldrich | P6148 |
| Bovine Serum Albumin | Sigma-Aldrich | A7906 |
| Triton X-100 | Sigma-Aldrich | 9002-93-1 |
| Sodium Cacodylate buffer | Electron Microscopy Sciences | 11654 |
| Tris | Fisher | BP152-5 |
| Ethylenediaminetetraacetic acid | Sigma-Aldrich | 6381-92-6 |
| Sodium Chloride | Sigma-Aldrich | 793566 |
| β-mercaptoethanol | Sigma-Aldrich | M6250 |
| Sodium dodecyl sulfate | Sigma-Aldrich | L3771 |
| Glycine | Fisher | BP381-5 |
| Sodium Citrate | Sigma-Aldrich | S-4641 |
| Metafectene Pro | Biontex | T040 |
| Critical Commercial Assays |  |  |
| Click-iT EdU imaging kit | Thermo Fisher Scientific | C10337 |
| SuperSignal™ West Femto Maximum Sensitivity Substrate | Thermo Fisher Scientific | 34095 |
| Duolink In Situ green kit Mouse/ Rabbit | Sigma Aldrich | DUO92014 |
| PureLink Genomic DNA mini kit | Thermo Fisher Scientific | K182001 |
| PowerUp SYBR Green Master Mix | Thermo Fisher Scientific | A25742 |
| Experimental Models: Cell Lines |  |  |
| Primary human fibroblasts (controls and DRP1 mutants) | Skin biopsies (Research Ethics Board of the Children's Hospital of Eastern Ontario (DRP1 mutants)) |  |
| Mouse Embryonic Fibroblasts | Dr. Luca Scorrano, University of Padua |  |
| U2OS | Gift from Dr. Edward Fon, McGill University |  |
| Oligonucleotides |  |  |
| Negative siRNA | Thermo Fisher Scientific | 4390843 |
| DRP1 siRNA (Mouse) | Thermo Fisher Scientific | 4390771 |
| DRP1 siRNA (Human) | Horizondiscovery | M-012092-01-0005 |
| CLIMP siRNA | Horizondiscovery | M-012755-02-0005 |
| SYNJ2BP siRNA | Horizondiscovery | M-021176-01-0005 |
| Recombinant DNA |  |  |
| mCherry | Addgene | 54517 |
| mCherry-tagged CLIMP63 | Addgene | 136293 |
| mcherry-FIS1 | gift from Uri Manor |  |
| mch-SYNJ2BP | gift from Marc Tramier |  |
| DRP1 K38A | Addgene | 45161 |
| Software and Algorithms |  |  |
| ImageJ | National Institutes of Health | https://imagej.nih.gov/ij/ |
| R | www.r-project.org/ | R4.0.5 |
| Imagelab | BioRad |  |
| Bitplane Imaris v. 9.5 | Oxford Instruments |  |
| Dragonfly | Object research systems |  |
| R studio | http://www.rstudio.com/ | Version 2023.03.0+386 |

