## Supplementary figures for "Contact sites between endoplasmic reticulum sheets and mitochondria regulate mitochondrial DNA replication and segregation"

Supplementary Figure 1

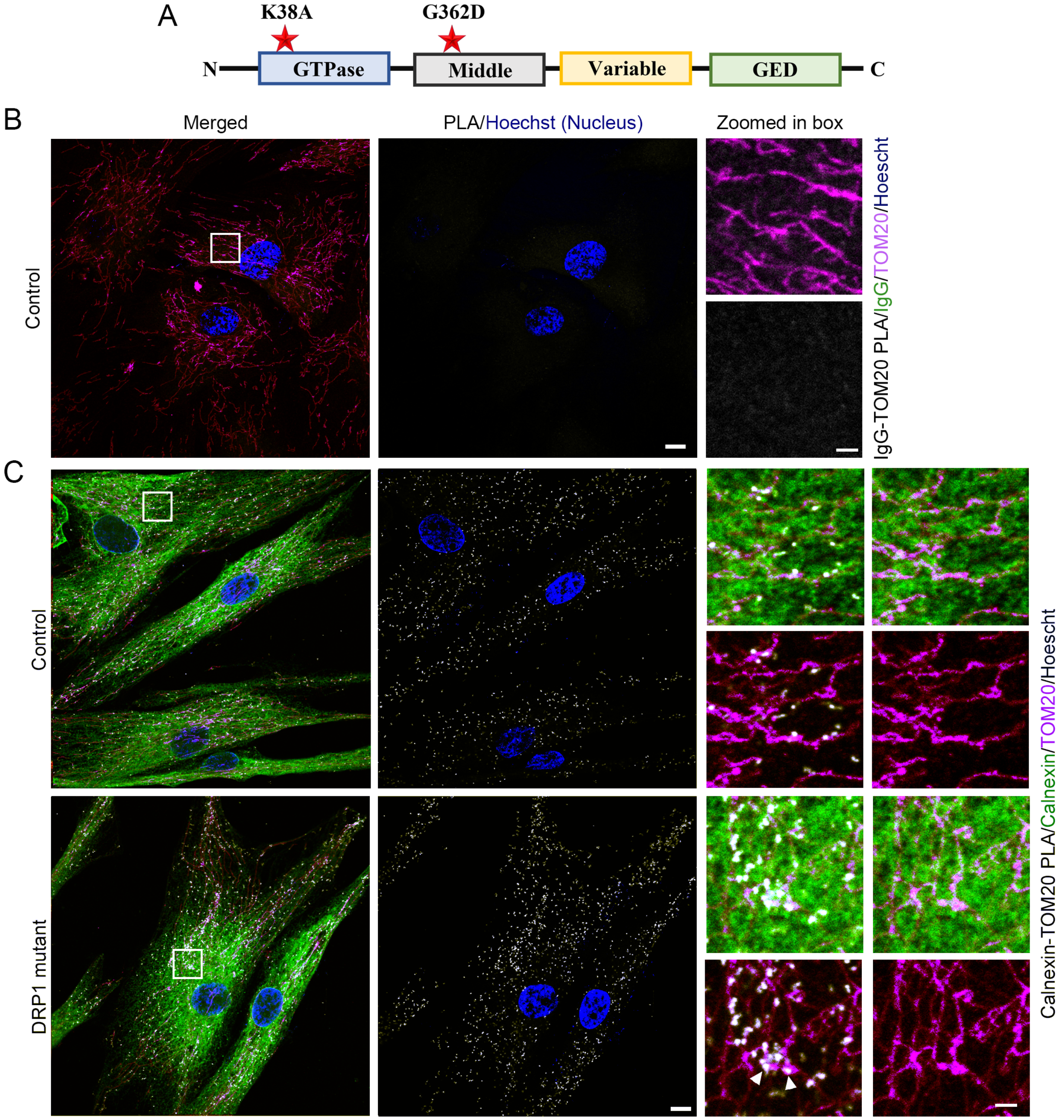


**Figure S1. Calnexin-TOM20 PLA** (related to Figure 2). (A) Schematic representation of DRP1 with the mutants that were used in this study labeled with a star. (B) Representative image of PLA (IgG and TOM20; white) for the antibody control (IgG), along with IgG (Green), TOM20 (mitochondria, Magenta) and nuclei (Hoechst, Blue) in control fibroblasts. (C) Representative images of control and DRP1 mutant fibroblasts showing the PLA for Calnexin and TOM20 (white), along with Calnexin (ER, green), TOM20 (mitochondria, magenta) and nuclei (Hoechst, Blue). Scale bar 10 μm. Scale bar for the enlarged areas 2 μm

Figure S2

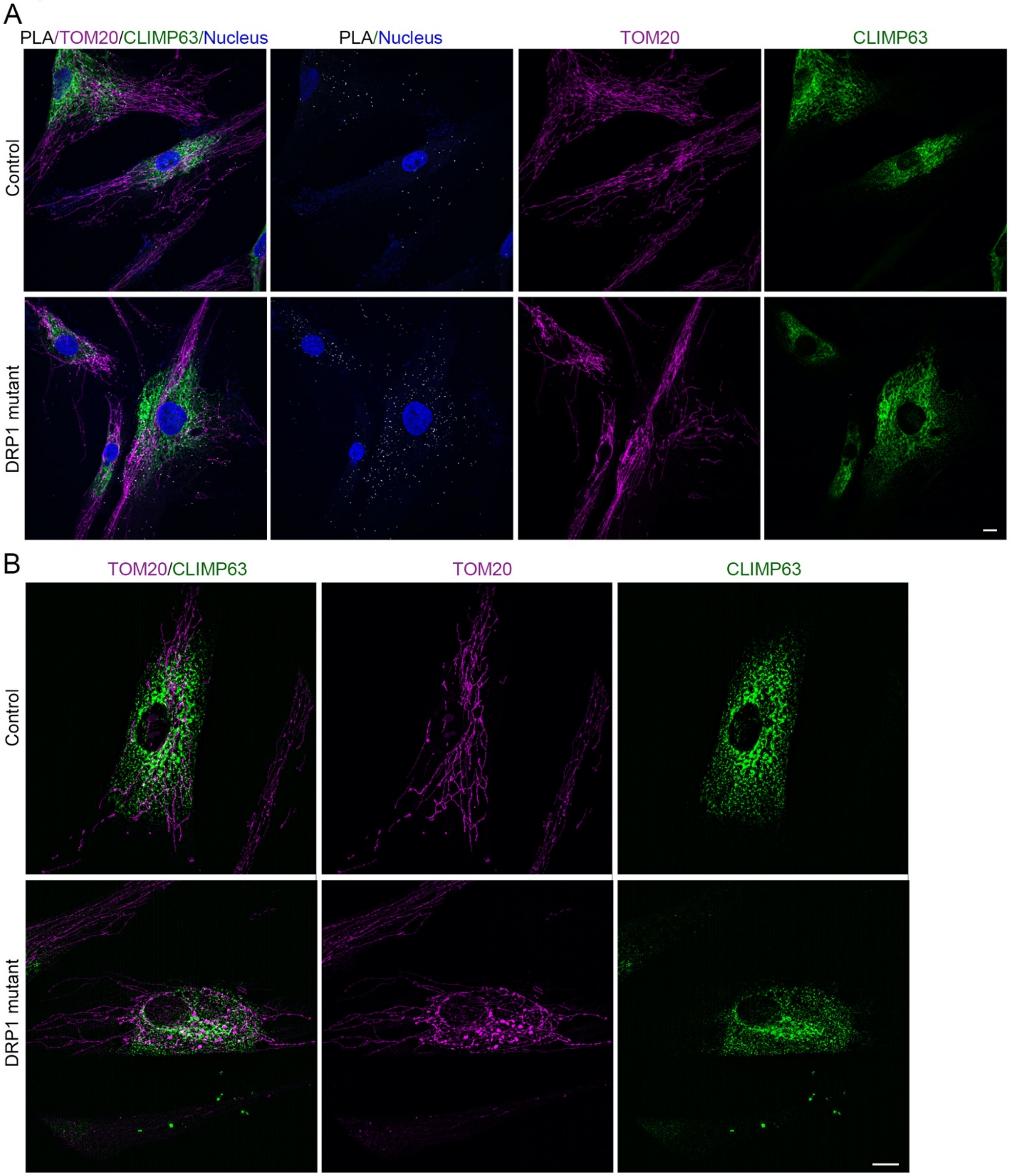


**Figure S2. CLIMP63-TOM20 contact sites in DRP1 mutant cells** (related to Figure 3). (A) Representative images of control and DRP1 mutant fibroblasts showing the PLA for CLIMP63 and TOM20 (white), along with CLIMP63 (ER, green), TOM20 (mitochondria, magenta) and nuclei (Hoechst, Blue). (B) Representative SIM images of control and DRP1 mutant fibroblasts stained for CLIMP63 (ER sheets, green) and Mitotracker orange (mitochondria, magenta). Scale bars 10 μm.

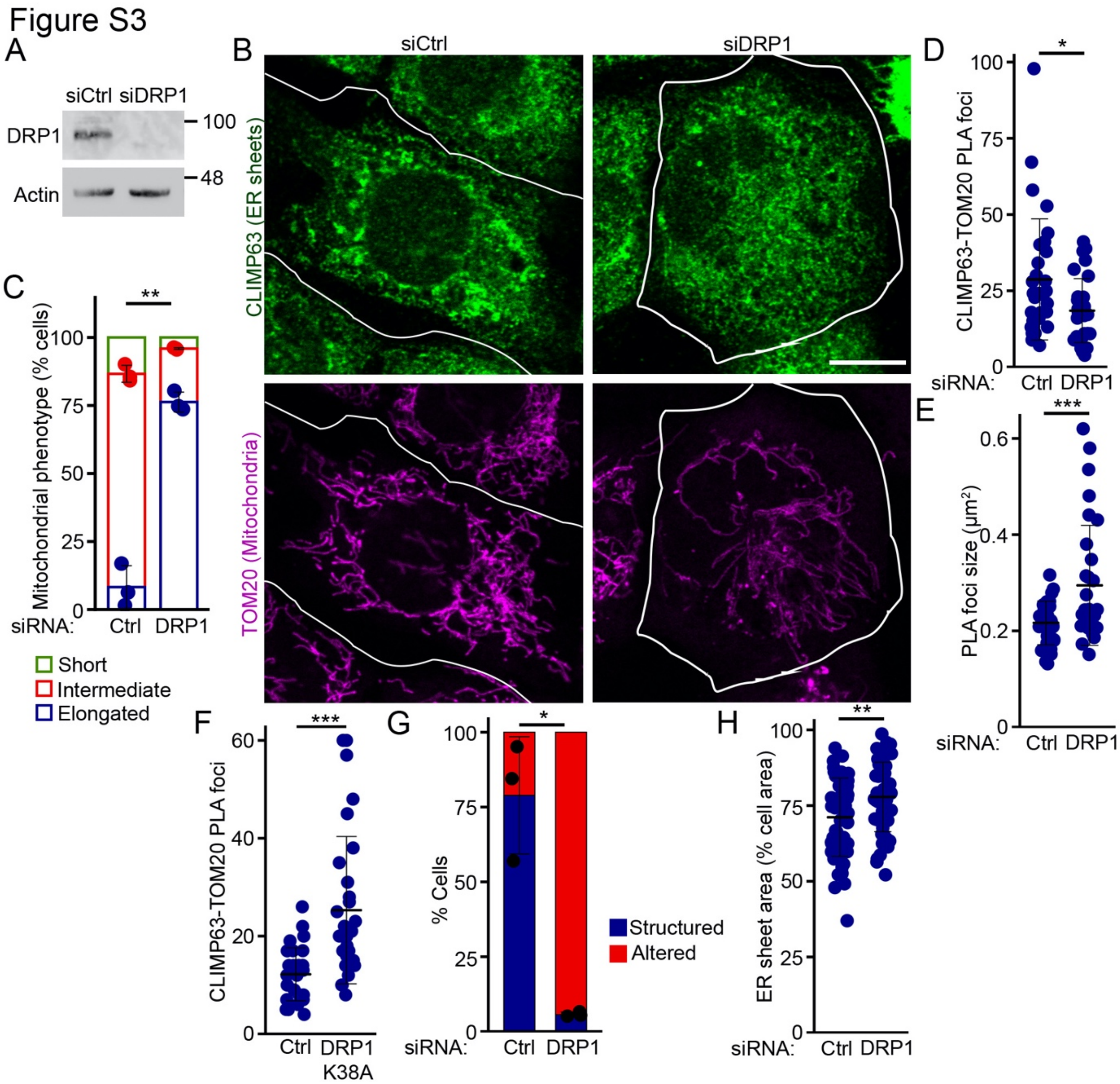


**Figure S3. Alterations in ER structure and mitochondrial contact sites following DRP1 knockdown in MEFs** (related to Figure 3). (A) WB showing DRP1 expression in WT MEFs transfected with a control siRNA (siCtrl) or a siRNA against DRP1 (siDRP1). (B) Representative images showing TOM20 (mitochondria) and CLIMP63 (ER sheets) staining in MEFs transfected with control and DRP1 siRNAs. White lines represent the cell edge. Scale bar 10 μm (C) Quantification of mitochondrial phenotypes (short (green), intermediate (red), elongated/hyperfused (blue)) in siCtrl and siDRP1 MEFs. Each point represents an independent experiment. Bars show the average ± SD. ** $p<0.01$ two-sided t-test (D-E) Quantification of CLIMP63-TOM20 PLA foci number (D) and size (E) in siCtrl and siDRP1 MEFs. Each data point represents one cell. Bars represent the average of 30 cells/condition in 3 independent experiments ± SD * $p<0.05$, *** $p<0.001$ two-sided t-test. (F) Quantification of CLIMP63-TOM20 PLA foci in MEFs transfected with DRP1K38A-HA and mCherry-Fis1 (to label transfected cells). Bars represent the average of 30 cells/condition in 3 independent experiments ± SD *** $p<0.001$ two-sided t-test. (G) Quantification of ER sheet structure as Structured (Blue) or Altered (red; presence of punctate structures and thick ER sheet patches) in WT MEFs transfected with a control siRNA (siCtrl) or a siRNA against DRP1 (siDRP1). Each point represents one independent experiment, with at least 20 cells quantified per experiment. Bars show the average ± SD. * $p<0.05$, Two-sided t-test using the data for Structured ER. (H) Quantification of ER sheet surface area relative to the total cell area (determined by DIC) in WT MEFs transfected with a control siRNA (siCtrl) or a siRNA against DRP1 (siDRP1). Each data point represents one cell. Bars represent the average of 54 cells in 3 independent experiments ± SD. ** $p<0.01$ two-sided t-test.

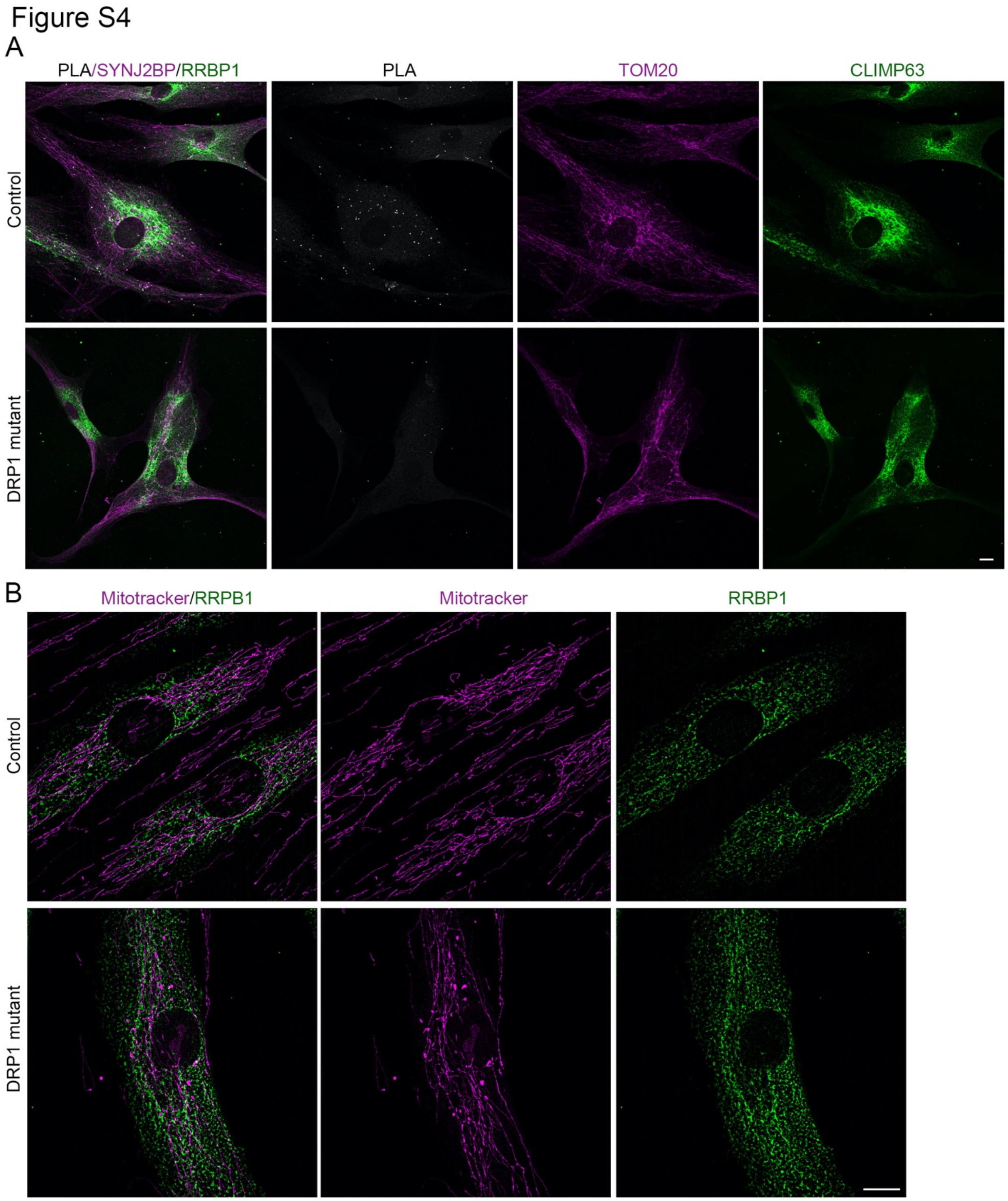


**Figure S4. RRBP1-SYNJ2BP contact sites in DRP1 mutant cells** (related to Figure 4). (A) Representative images of control and DRP1 mutant fibroblasts showing the PLA for RRBP1 and SYNJ2BP (white), along with RRBP1 (ER, green) and SYNJ2BP (mitochondria, magenta) (Hoechst, Blue). (B) Representative SIM images of control and DRP1 mutant fibroblasts stained for RRBP1 (ER sheets, green) and Mitotracker orange (mitochondria, magenta). Scale bars 10 µm.

Figure S5

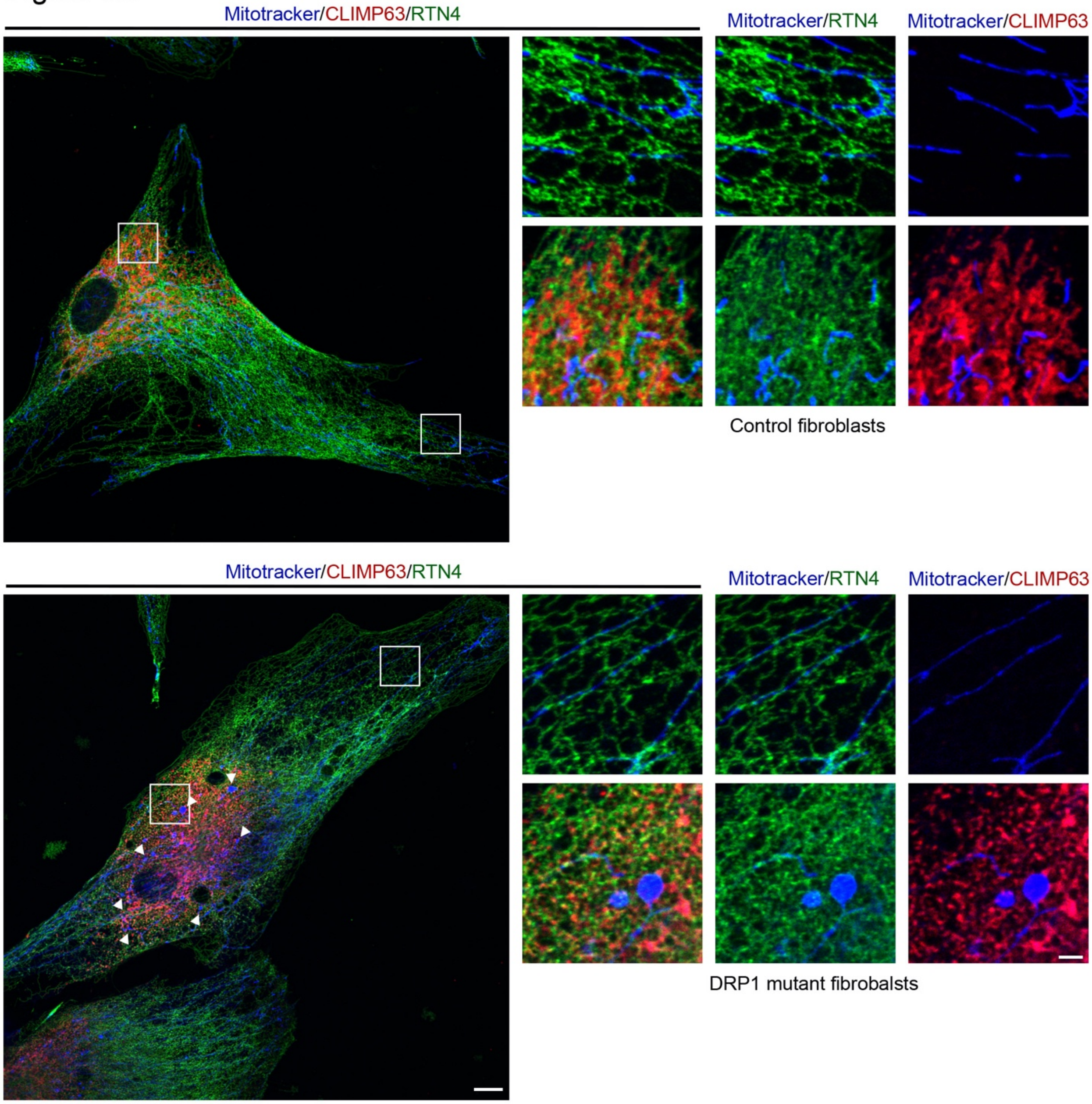


**Figure S5. ER structure in DRP1 mutant fibroblasts** (related to Figure 6). Representative images of control (Top) and DRP1 mutant (Bottom) human fibroblasts showing CLIMP63 (ER sheets, Red), RTN4 (ER, Green) and mitochondria (Mitotracker orange, Blue). The enlarged boxed areas show ER structure in the perinuclear area (CLIMP63-positive) and periphery of the cells. Scale bar 10 µm, 2 µm for the enlarged images. Arrowheads denote the localisation of mitobulbs within CLIMP63-positive perinuclear area.

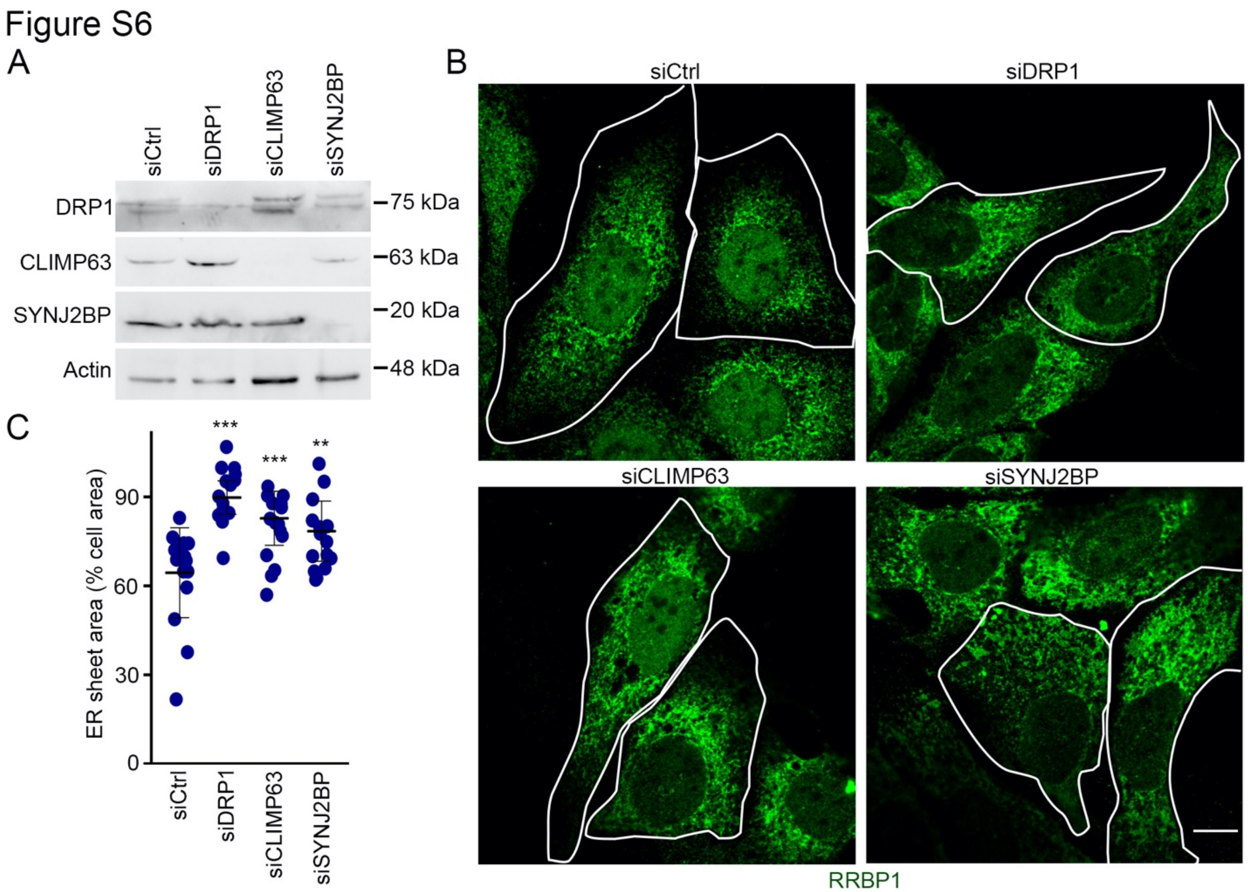


**Figure S6. ER sheet alterations in U2OS cells knocked down for DRP1, CLIMP63 or SYNJ2BP** (related to Figure 7). (A) WB showing the expression of the indicated proteins in U2OS cells transfected with either a control siRNA (siCtrl), a siRNA against DRP1 (siDRP1), CLIMP63 (siCLIMP63) or SYNJ2BP (siSYNJ2B). (B) Representative images showing RRBP1 (ER sheets) staining in U2OS transfected with the indicated siRNAs. White lines represent the cell edge. Scale bar 10 µm. (C) Quantification of ER sheet surface area relative to the total cell area (determined by DIC). Each data point represents one cell. Bars represent the average of 15 cells in 3 independent experiments ± SD. One-way ANOVA *** $p<0.001$, ** $p<0.01$.

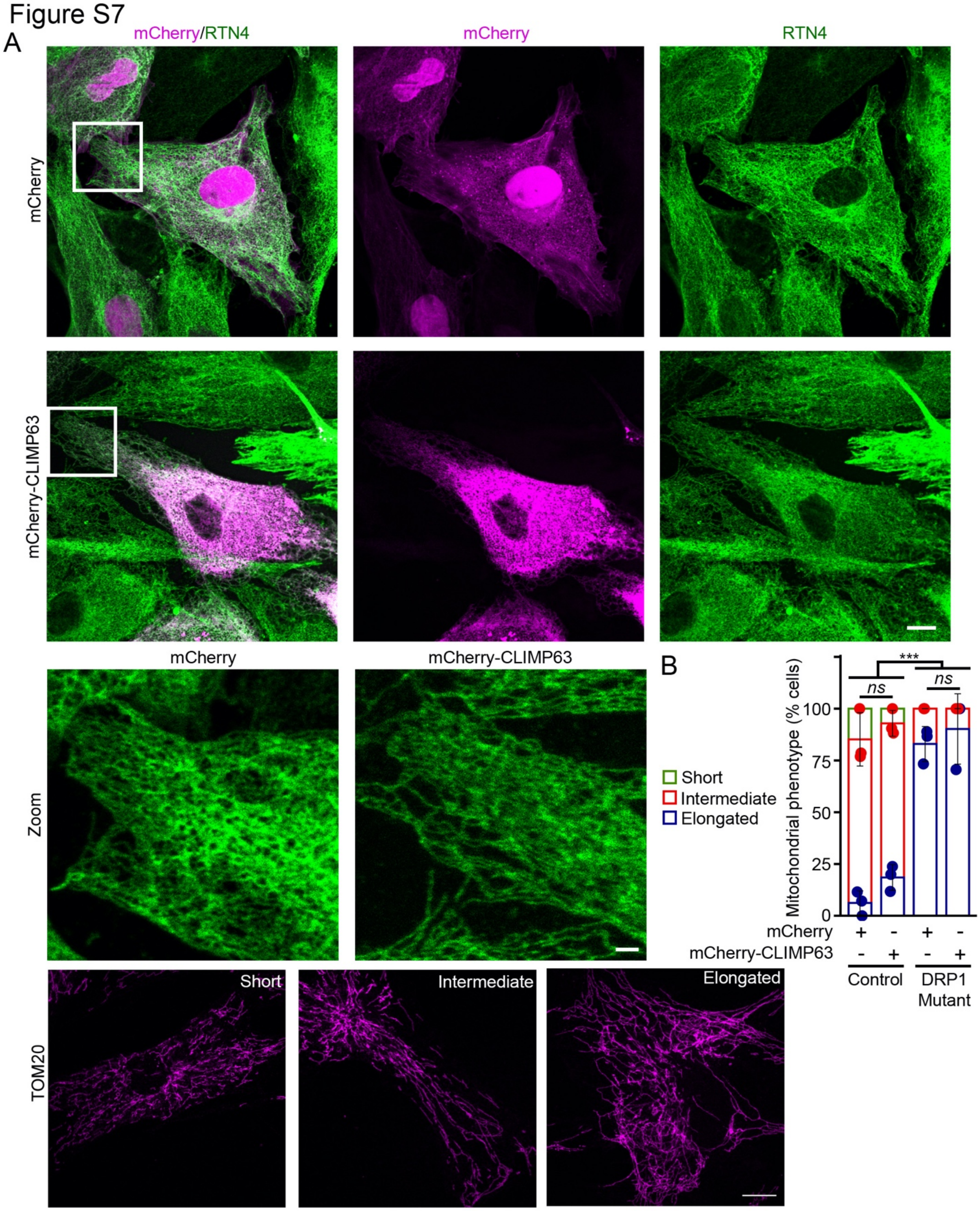


**Figure S7. ER tubules are not affected by CLIMP63 expression** (related to Figure 8). (A) Representative images of Control cells expressing mCherry and mCherry-CLIMP63 (Magenta) and immunolabelled for RTN4 (ER, green). Scale bar 10 µm, 2 µm for the zoomed images. (B)

Quantification of mitochondrial phenotypes (short (green), intermediate (red), elongated/hyperfused (blue)) in mCherry and mCherry-CLIMP63 expressing control and DRP1 mutant fibroblasts. Each point represents an independent experiment. Bars show the average ± SD. Two-way ANOVA. *** $p<0.001$, ns not significant.

Figure S8

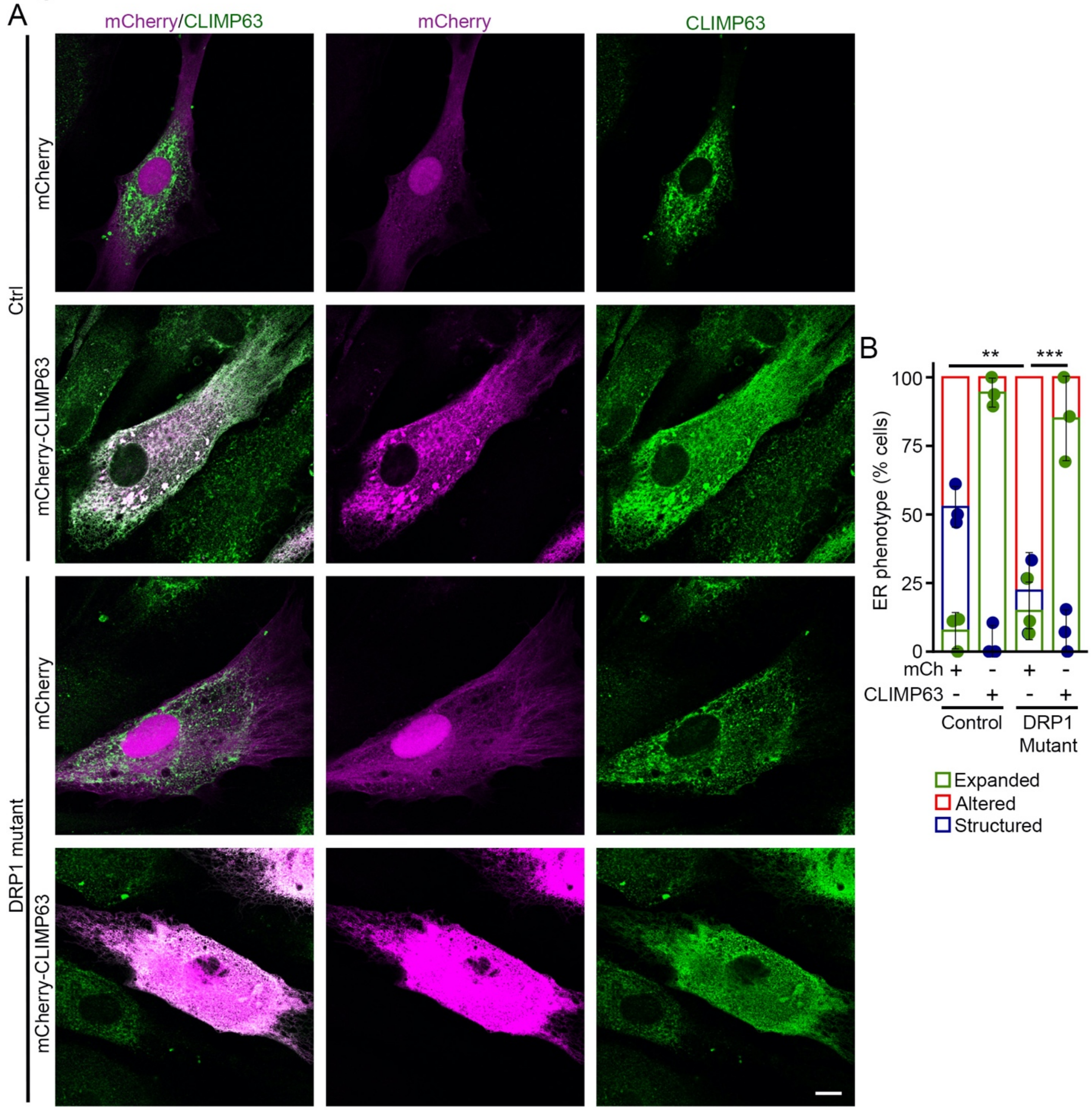


**Figure S8. CLIMP63 expression alters ER sheet structure in control and DRP1 mutants** (related to Figure 8). (A) Representative images of control and DRP1 mutant fibroblasts expressing mCherry or mCherry-CLIMP63 (Magenta) and immunolabelled for CLIMP63 (ER sheets, green). Scale bar 10 µm. (B) Quantification of ER sheet structure from images in (A) as Structured (Blue), Altered (red; presence of punctate structures and thick ER sheet patches) or expanded (a large cell area covered by structured ER sheets). Each point represents one independent experiment, with at least 10 transfected cells quantified per experiment.

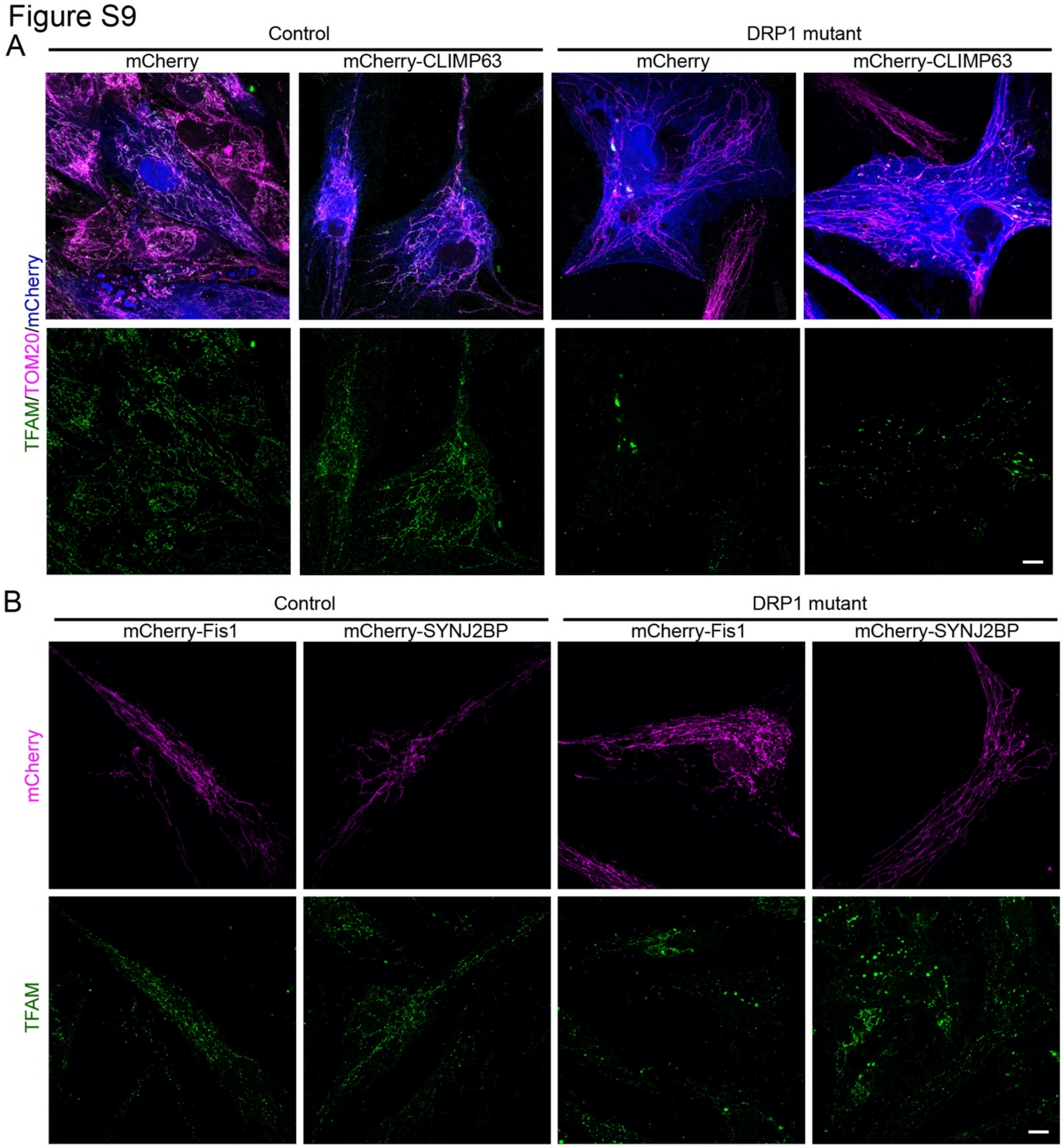


**Figure S9. CLIMP63 expression rescues nucleoid aggregation in DRP1 mutants** (related to figure 8). (A) Representative images of control and DRP1 mutant fibroblasts expressing mCherry or mCherry-CLIMP63 (blue) immunolabeled for TFAM (nucleoids, Green) and TOM20 (mitochondria, Magenta). (B) Representative images of control and DRP1 mutant fibroblasts expressing mCherry-Fis1 or mCherry-SYNJ2BP (Magenta) immunolabeled for TFAM (nucleoids, Green). Scale bars 10 µm.

Figure S10

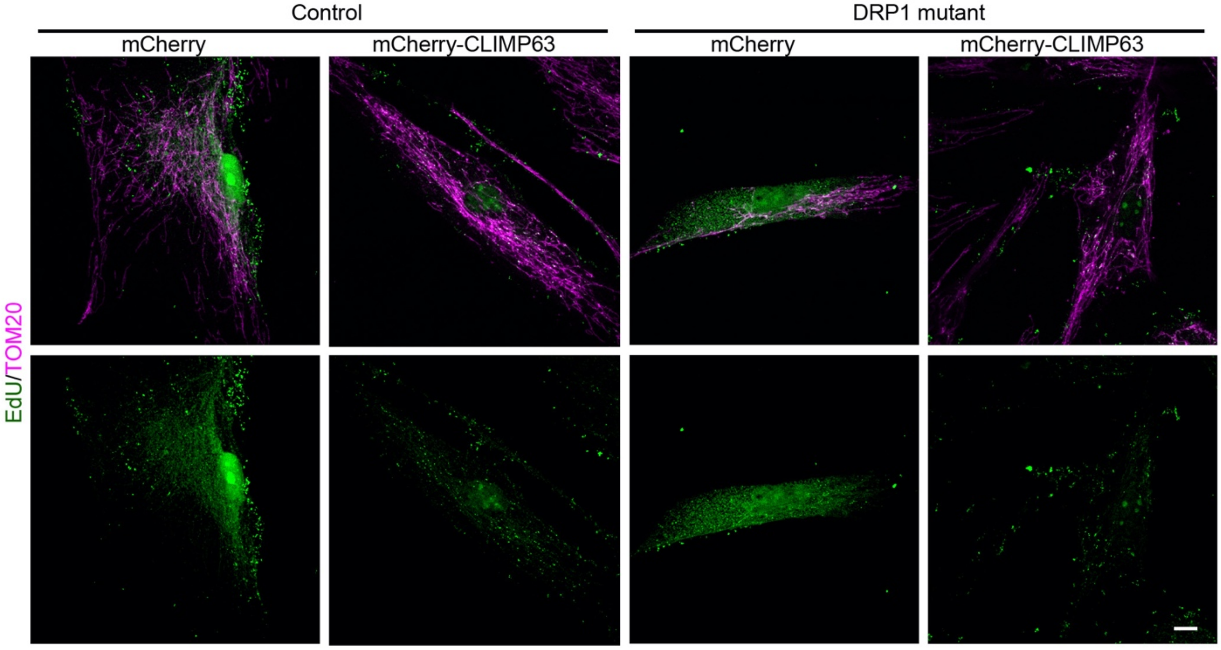


**Figure S10. CLIMP63 expression rescues nucleoid replication in DRP1 mutants** (related to figure 9). Representative images of control and DRP1 mutant fibroblasts expressing mCherry or mCherry-CLIMP63 immunolabeled for EdU (replicating nucleoids, Green) and TOM20 (mitochondria, Magenta). Scale bar 10 µm.
